# Methacrylate-Modified Gold Nanoparticles Enable Non-Invasive Monitoring of Photocrosslinked Hydrogel Scaffolds

**DOI:** 10.1101/2022.01.26.477960

**Authors:** Lan Li, Carmen J. Gil, Tyler A. Finamore, Connor J. Evans, Martin L. Tomov, Liqun Ning, Andrea Theus, Gabriella Kabboul, Vahid Serpooshan, Ryan K. Roeder

**Affiliations:** Department of Aerospace and Mechanical Engineering, Bioengineering Graduate Program, Materials Science and Engineering Graduate Program, University of Notre Dame, Notre Dame, IN 46556, USA; Notre Dame Center for Nanoscience and Technology (ND*nano*), University of Notre Dame, Notre Dame, IN 46556, USA; Wallace H. Coulter Department of Biomedical Engineering, Georgia Institute of Technology, Atlanta, GA 30332, USA; Department of Pediatrics, Emory University School of Medicine, Emory University, Atlanta, GA 30322, USA

**Keywords:** Computed Tomography, Gelatin Methacryloyl (GelMA) Hydrogel, Gold Nanoparticles, Photopolymerization, Tissue Engineering Scaffold

## Abstract

Photocrosslinked hydrogels, such as methacrylate-modified gelatin (gelMA) and hyaluronic acid (HAMA), are widely utilized as tissue engineering scaffolds and/or drug delivery vehicles, but lack a suitable means for non-invasive, longitudinal monitoring of surgical placement, biodegradation, and drug release. Therefore, we developed a novel photopolymerizable X-ray contrast agent, methacrylate-modified gold nanoparticles (AuMA NPs), to enable covalent-linking to methacrylate-modified hydrogels (gelMA and HAMA) in one-step during photocrosslinking and non-invasive monitoring by X-ray micro-computed tomography (micro-CT). Hydrogels exhibited a linear increase in X-ray attenuation with increased Au NP concentration to enable quantitative imaging by contrast-enhanced micro-CT. The enzymatic and hydrolytic degradation kinetics of gelMA-Au NP hydrogels were longitudinally monitored by micro-CT for up to one month *in vitro*, yielding results that were consistent with concurrent measurements by optical spectroscopy and gravimetric analysis. Importantly, AuMA NPs did not disrupt the hydrogel network, rheology, mechanical properties, and hydrolytic stability compared with gelMA alone. GelMA-Au NP hydrogels were thus able to be bioprinted into well-defined three-dimensional architectures supporting endothelial cell viability and growth. Overall, AuMA NPs enabled the preparation of both conventional photopolymerized hydrogels and bioprinted scaffolds with tunable X-ray contrast for noninvasive, longitudinal monitoring of placement, degradation, and NP release by micro-CT.

## 1. Introduction

Photocrosslinked hydrogels, such as methacrylate-modified gelatin (gelMA),^1–3^ hyaluronic acid (HAMA),^4–6^ and collagen (colMA),^7–9^ are widely utilized as tissue engineering scaffolds and drug delivery vehicles due to enabling precision manufacturing (e.g., 3D printing) of biodegradable materials with tunable properties, and the incorporation of sensitive cells and/or biomolecules.^10 12^ However, there is currently no established means for noninvasive, longitudinal, and volumetric monitoring of hydrogel scaffolds once implanted *in vivo.* Conventional methods to longitudinally monitor scaffold degradation and integration, such as histology and mechanical testing, require invasive excision of multiple tissue samples.^13^ Therefore, various imaging modalities have been investigated for noninvasive assessment of hydrogel degradation and drug delivery,^13,14^ including ultrasound elasticity imaging,^15,16^ photoacoustic imaging,^17,18^ near-infrared fluorescence imaging,^19–24^ magnetic resonance imaging,^24–27^ and X-ray computed tomography (CT).^28–30^ Among these, CT imaging is advantageous in providing low cost, three-dimensional (3D), deep tissue imaging at high spatial and temporal resolution, but is limited by low soft tissue contrast.

CT imaging contrast is derived from the X-ray attenuation of the scaffold material relative to adjacent tissue.^31^ Hydrogels and other soft biomaterials exhibit similar X-ray attenuation to soft tissues and thus low X-ray contrast.^13^ Therefore, contrast agents are required to overcome this limitation. Gold nanoparticles (Au NPs) have become the most widely utilized X-ray contrast agent in preclinical research due to exhibiting strong X-ray attenuation while facilitating facile synthesis and surface modification.^31,32^ Therefore, the incorporation of Au NPs within photocrosslinked hydrogels could enable non-invasive, post-operative imaging of surgical placement and longitudinal, quantitative imaging of degradation and/or drug delivery. However, the method by which Au NPs are integrated into a hydrogel is crucial for achieving the desired functionality.^33–35^

Au NPs have been incorporated within hydrogels by physical and chemical means.^33–35^ In physical incorporation, Au NPs are mixed into the prepolymer solution and entrapped within the hydrogel during crosslinking.^36–38^ Physical incorporation is simple and flexible but may suffer from disrupting the hydrogel network and properties, including premature or uncontrolled (burst) release of NPs which limits the effective time for longitudinally monitoring hydrogel function.^36–39^ In chemical incorporation, Au NPs are surface functionalized with ligands that are able to be chemically-coupled to hydrogel macromolecules.^29,35,40,41^ Chemically-incorporated NPs are immobilized such that their release coincides with hydrolytic or enzymatic degradation of the hydrogel for accurate and reliable monitoring.^29^ However, the chemical incorporation of NPs in hydrogels typically requires modification of both NP surfaces and hydrogel macromolecules, involving multi-step reactions with potentially undesirable side reactions, prior to photocrosslinking the hydrogel.^40–43^ Thus, simple and flexible methods are needed for the chemical incorporation of Au NPs in photocrosslinked hydrogels with minimal disruption of the hydrogel structure and properties.

Therefore, the objective of this study was to investigate a photopolymerizable X-ray contrast agent, methacrylate-modified Au NPs (AuMA NPs), which can be directly conjugated to methacrylate-modified hydrogels (gelMA and HAMA) in one-step during photocrosslinking to subsequently enable non-invasive, longitudinal monitoring of scaffold degradation and/or drug delivery. GelMA-Au NP hydrogels prepared by one-step photopolymerization (1-step gelMA-Au) were compared with gelMA alone, gelMA with physically-entrapped Au NPs (gelMA+Au), and gelMA chemically-coupled with Au NPs using carbodiimide/succinimide (EDC/NHS) chemistry prior to photocrosslinking (2-step gelMA-Au) (Fig. 1). The effective swelling ratio, mechanical properties and degradability of photocrosslinked gelMA-Au NP hydrogels and the rheological properties of bioinks were characterized and compared with normal gelMA. The hydrolytic and enzymatic degradation kinetics of gelMA-Au NP hydrogels were measured longitudinally by contrast-enhanced micro-CT, and compared with concurrent measurements by optical spectroscopy and gravimetric analysis. In addition, gelMA-Au NP bioinks were printed into 3D architectures and loaded with endothelial cells to evaluate cell viability.

**Fig. 1.**
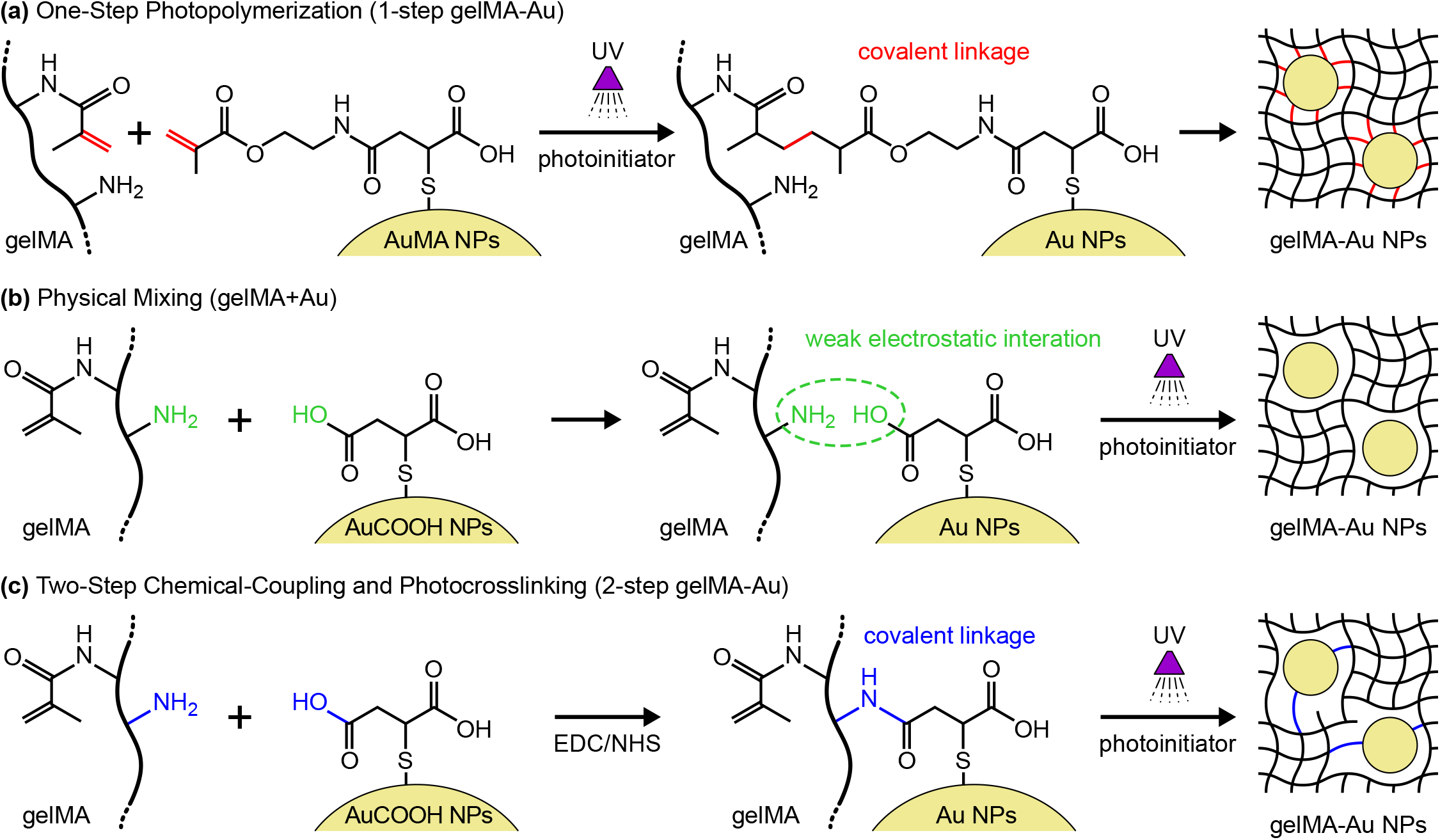
Schemes for the preparation of photopolymerized gelMA-Au NP hydrogels: (a) one-step photopolymerization of gelMA and AuMA NPs (1-step gelMA-Au) to form covalent linkages, (b) mixing gelMA and AuCOOH NPs (gelMA+Au) with weak electrostatic interactions to physically-entrap Au NPs within the gelMA network, and (c) two-step chemical-coupling and photocrosslinking (2-step gelMA-Au) where covalent linkages between gelMA macromolecules and AuCOOH NPs are formed by EDC/NHS chemistry prior to photocrosslinking gelMA.

## 2. Materials and methods

### 2.1. Synthesis of AuCOOH NPs

AuCOOH NPs were synthesized by surface functionalizing bare Au NPs, ~12 nm in diameter prepared by the citrate reduction method, with mercaptosuccinic acid (MSA).^29,44^ Briefly, 0.1 g gold (III) chloride trihydrate (HAuCl_4_·3H_2_O, 99.9%, Sigma-Aldrich) was added to 500 mL deionized (DI) water and heated to boiling while stirring. Once boiling, 0.5 g trisodium citrate dihydrate (C_6_H_5_Na_3_O_7_·2H_2_O, ACS reagent, Sigma-Aldrich) was added to the mixture. The mixture was boiled for another 20 min, cooled to room temperature and stirred overnight. As-prepared Au NPs were collected in a volumetric flask and titrated to 500 mL. An aqueous solution containing 15 mL of 10 mM MSA (C_4_H_6_O_4_S, 97%, Sigma-Aldrich) was added to the Au NP solution and stirred overnight. As-prepared AuCOOH NPs were collected by centrifugation (Sorvall RC-6 Plus, Thermo Scientific) at ~11,000*g* for 1 h and thrice washed with DI water. The final AuCOOH NP stock solution was concentrated to 94 mM as verified by inductively-coupled plasma optical emission spectroscopy (ICP-OES).

### 2.2. Synthesis of AuMA NPs

AuMA NPs were synthesized by covalently-linking AuCOOH NPs with 2-aminoethyl methacrylate (AEMA, 90%, C_6_H_11_NO_2_·HCl, Sigma-Aldrich) using carbodiimide/succinimide chemistry (Fig. 2a). First, 0.5 mmol AuCOOH NPs were added to 200 mL ethanol (80% v/v) containing 1.44 g 1-ethyl-3-(3-dimethylaminopropyl) carbodiimide hydrochloride (EDC, Sigma-Aldrich) and 0.65 g *N*-hydroxysulfosuccinimide (NHS, Sigma-Aldrich), which was then mixed with another 200 mL ethanol (80% v/v) containing 0.495 g fully dissolved AEMA, such that the molar ratio of Au:EDC:NHS:AEMA was 1:15:6:6. The mixture was vigorously stirred under nitrogen protection for 24 h at room temperature to obtain AuMA NPs. AuMA NPs were also prepared with a higher degree of methacrylation (Au:EDC:NHS:AEMA = 1:75:30:30) but exhibited strong hydrophobicity and were not utilized further. After the reaction, AuMA NPs were collected by centrifugation at 8400*g* for 30 min and washed thrice with DI water. The final AuMA NP stock solution was concentrated to 90 mM as verified by ICP-OES.

**Fig. 2.**
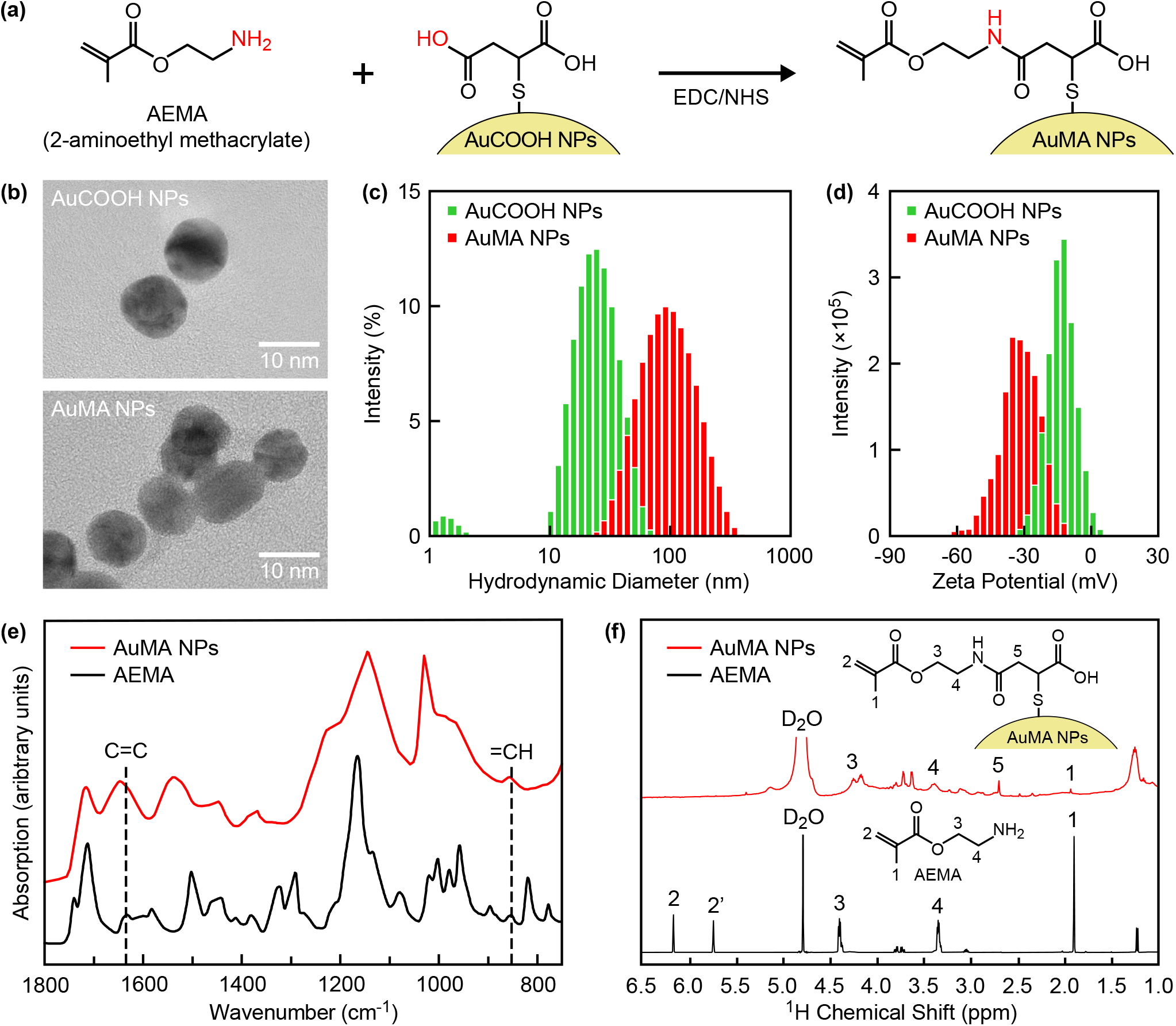
Synthesis and characterization of AuMA NPs. (a) Scheme for the preparation of AuMA NPs by covalently linking AEMA with AuCOOH NPs using EDC/NHS chemistry. (b) Representative TEM micrographs showing as-prepared AuCOOH and AuMA NPs, which were both spherical and well-dispersed, although AuMA NPs formed dispersible nanoclusters in aqueous solution. The (c) hydrodynamic diameter and (d) zeta potential distributions of as-prepared AuCOOH and AuMA NPs measured by DLS, which confirmed solubility and colloidal stability in aqueous solution. (e) FTIR and (f) ^1^H NMR spectra of as-prepared AuMA NPs compared with AEMA, which verified methacrylate surface modification.

### 2.3. Characterization of AuCOOH and AuMA NPs

#### 2.3.1. ICP-OES

The concentration of as-prepared AuCOOH NPs and AuMA NPs, as well as the concentration of Au NPs released from hydrogels into supernatant media during *in vitro* hydrolytic and enzymatic degradation, was measured using ICP-OES. As-prepared or supernatant Au NP solutions were digested in 5% aqua regia before analysis. Calibration curves were established by diluting certified standard Au solutions (Assurance Grade, SPEX CertiPrep).

#### 2.3.2. Transmission electron microscopy (TEM)

The size and morphology of as-prepared AuCOOH and AuMA NPs were characterized by TEM (JEOL 2011, JEOL) at an accelerating voltage of 120 kV. Specimens were prepared by pipetting 10 μL of as-prepared NPs onto carbon-coated grids and evaporating the solvent in an oven at 60°C. The mean (± standard deviation) diameter and size distribution were measured from at least 100 NPs sampled from digital images at 120,000X magnification (ImageJ v1.51).

#### 2.3.3. Dynamic light scattering (DLS)

The hydrodynamic particle diameter and zeta potential were measured by DLS (Zetasizer ZS90, Malvern Instruments) at 25°C. Specimens were prepared by dispersing NPs in DI water at 0.5 mM and 5 mM concentration for hydrodynamic diameter and zeta potential measurements, respectively. The mean (± standard deviation) hydrodynamic diameter and zeta potential were measured from three sample replicates.

#### 2.3.4. Fourier transform infrared spectroscopy (FTIR)

Methacrylate surface functionalization of as-prepared AuMA NPs was verified by FTIR transmission spectra of AEMA powder and lyophilized AuMA NPs acquired using an attenuated total reflection (ATR) module (Tenor 27, Bruker) as an average of 64 scans over 4000–400 cm^−1^ at room temperature.

#### 2.3.5. Nuclear magnetic resonance (NMR)

Methacrylate surface functionalization of as-prepared AuMA NPs was also characterized by ^1^H NMR spectra of AEMA powder and lyophilized AuMA NPs acquired at 25°C on a spectrometer (AVANCE III HD 500, Bruker) with 8012.8 Hz spectral width, 4.0 s acquisition time, 6.5 ms pulse, and referenced to the solvent peak of deuterium oxide (D2O) at 4.79 ppm. Specimens were prepared by dissolving AEMA powder and lyophilized AuMA NPs in 0.6 mL of D2O (Sigma-Aldrich) at a concentration of 10 and 30 mg/mL, respectively.

### 2.4. Preparation and characterization of gelMA and gelMA-Au NP prepolymer solutions

#### 2.4.1. GelMA

GelMA with a degree of functionalization (DoF) >75-80% was prepared using previously established methods.^45,46^ Briefly, 8 g porcine gelatin (Sigma) was dissolved in 100 mL phosphate-buffered saline (PBS) at 50°C under stirring, followed by the dropwise addition of 8 mL methacrylic anhydride (MAA, Sigma) which was maintained at 50°C under stirring for 3 h. The gelMA solution was then centrifuged to remove impurities, diluted with 100-200 mL PBS, and dialyzed for 7 d to remove unreacted MAA, lyophilized, and stored away from light at −°C until further use. Lyophilized gelMA powder was reconstituted in PBS at 40% w/v and heating to 60°C for the preparation of prepolymer solutions.

#### 2.4.2. GelMA and gelMA-Au NP prepolymer solutions

All gelMA and gelMA-Au NP prepolymer solutions were prepared with 10 or 20% w/v gelMA and up to 37 mM Au NPs. Prepolymer solutions for one-step photopolymerized gelMA-Au NP hydrogels (1-step gelMA-Au) and physically-mixed gelMA+Au NP hydrogels (gelMA+Au) were prepared by mixing appropriate volumes of the gelMA prepolymer solution with AuMA or AuCOOH NPs, respectively, at 60°C and vortexing for 2 min. Prepolymer solutions for gelMA-Au NP hydrogels prepared by two-step chemical-coupling and photocrosslinking (2-step gelMA-Au) were prepared by covalently-linking gelMA and AuCOOH NPs via EDC/NHS chemistry prior to photocrosslinking. Appropriate volumes of the gelMA prepolymer solution were mixed with AuCOOH NPs at 60°C and vortexed for 2 min. The mixture was loaded into a 4.78 mm inner diameter mold, cooled to 4°C to induce thermal gelation, removed from the mold, and sectioned into cylindrical specimens, 5.0 mm in height, using a razor blade. Specimens were then soaked in 30 mL ethanol (80% v/v) containing EDC and NHS at 4°C with a Au:EDC:NHS molar ratio of 1:1:1 for 18 h. A higher Au:EDC:NHS molar ratio (Au:EDC:NHS = 1:160:64 for 6 h) was also investigated but compromised the thermal sol-gel transition and was not utilized further. After the reaction, specimens were thrice washed with DI water and heated above 60°C to obtain a chemically-coupled gelMA-Au NP prepolymer solution.

#### 2.4.3. Rheological properties

GelMA and gelMA-Au NP prepolymer solutions comprising 20% w/v gelMA and 10 mM Au NPs were characterized by loading 700 μL into the parallel plates (8 mm diameter, 1000 μm gap) of a rheometer (Discovery HR-2, TA Instruments). Storage moduli (*G*’) and loss moduli (*G*”) were measured at a constant frequency (1 Hz) and strain amplitude (1 %) over a temperature sweep from 37 to 5°C at a cooling rate of 3°C/min. The flow behavior of prepolymer solutions was also measured over a steady state strain rate sweep from 0.1 to 100 s ^1^ at 25°C. Three sample replicates were measured for each test and group.

### 2.5. Preparation and characterization of gelMA and gelMA-Au NP hydrogels

#### 2.5.1. Photocrosslinking

GelMA and gelMA-Au NP prepolymer solutions comprising 10 or 20% w/v gelMA and up to 37 mM Au NPs were supplemented with 0.5-1.0% w/v Irgacure 2959 (Sigma-Aldrich) or lithium phenyl-2,4,6-trimethylbenzoylphosphinate (LAP, Allevi) photoinitiator and incubated for 1 h, 24 h, or 7 d at 4°C. Prepolymer solutions with photoinitiator were loaded into cylindrical molds (4.78 mm inner diameter, 3 mm height) and photocrosslinked under an ultraviolet (UV) light source (OmniCure S1500, 320-390 nm) at 7, 15 or 30 mW/cm^2^ for 4-6 min at ambient temperature. After initial investigations to optimize the above photocrosslinking parameters, all gelMA and gelMA-Au NP hydrogels were prepared with 20% w/v gelMA and up to 37 mM Au NPs by incubating prepolymer solutions with 0.5% w/v LAP photoinitiator for 24 h at 4°C and photocrosslinking under UV irradiation at 30 mW/cm^2^ for 4 min.

#### 2.5.2. Contrast-enhanced micro-CT

The X-ray attenuation of gelMA-Au NP hydrogels was measured by micro-CT (μCT-80, Scanco Medical AG) for as-prepared gelMA-Au NP hydrogels comprising 20% w/v gelMA and 0-37 mM Au NPs. PBS and rat myocardium were imaged as soft tissue controls. Micro-CT images were also acquired for longitudinal measurements of degradation in gelMA-Au NP hydrogels initially containing 10 mM Au NPs and 3D bioprinted gelMA-Au NP constructs. Micro-CT images were acquired at 70 kVp tube potential, 144 μA beam current, and 0.5 mm Al beam filtration with 125 projections at 800 ms integration time. Noise in grayscale images was reduced using a Gaussian filter (sigma = 0.8, support = 1). 3D images were reconstructed with a 100 μm isotropic voxel size chosen to replicate *in vivo* imaging methods. The linear X-ray attenuation (cm^−1^) was measured within a volume of interest (VOI) and converted to Hounsfield units (HU) using internal calibration to air and water as −1000 and 0 HU, respectively. The VOI included the entire hydrogel volume for as-prepared gelMA-Au NP hydrogels or the entire volume of the media within Eppendorf tubes for longitudinal measurements of gelMA-Au NP hydrogel degradation. The total volume of the hydrogel and/or hydrogel fragments was segmented from the media using a fixed global threshold of 64, which corresponded to a linear attenuation of 0.51 cm^−1^ or 61 HU. Three sample replicates were imaged for each Au NP concentration in as-prepared gelMA-Au NP hydrogels. Five sample replicates were imaged longitudinally for each group during degradation of gelMA-Au NP hydrogels.

#### 2.5.3. Swelling ratio

The effective swelling ratio of gelMA and gelMA-Au NP hydrogels comprising 20% w/v gelMA and 10 mM Au NPs was measured by gravimetric analysis.^3,47,48^ Hydrogels were incubated in Dulbecco’s PBS (DPBS, Sigma-Aldrich) at 37°C for 24 h to reach the equilibrium swelling. The effective swelling ratio was measured by the total mass change after 24 h equilibrium relative to the initial mass. Ten sample replicates were measured for each group.

#### 2.5.4. Compression testing

GelMA and gelMA-Au NP hydrogels comprising 20% w/v gelMA and 10 mM Au NPs were loaded in unconfined uniaxial compression to 80% strain at a displacement rate of 0.05 mm/s using an electromagnetic test instrument (ElectroForce 3220, Bose). Hydrogels were incubated in DPBS for 24 h to reach equilibrium swelling prior to testing; specimens measured ~4 mm in diameter and ~3 mm in height. Load and displacement data were collected at a sampling rate of 10 Hz using a 25 lb load cell (MBP, Interface) and linear variable displacement transducer, respectively. The compressive modulus was calculated as the slope of the stress–strain curve over 10-20% strain using linear least squares regression (*R*^2^ > 0.95). At least three and at most five sample replicates were tested for each group.

### 2.6. In vitro hydrolytic and enzymatic degradation of gelMA and gelMA-Au NP hydrogels

GelMA and gelMA-Au NP hydrogels comprising 20% w/v gelMA and 10 mM Au NPs were evaluated using *in vitro* models of hydrolytic and enzymatic degradation.^29^ Hydrogels were incubated in DPBS for 24 h prior to degradation to establish the day 0 time point. Hydrogels were subsequently placed in Eppendorf tubes containing 0.2 mL DPBS for hydrolysis or 0.02 mg/mL bacterial (*Clostridium histolyticum*) collagenase (Collagenase B, Roche) in 0.2 mL DPBS for enzymolysis, and incubated at 37°C. Hydrolytic and enzymatic degradation were measured longitudinally at 24 h time points for up to one month. At each time point, the supernatant media was separated from hydrogels and/or hydrogel fragments by centrifugation at 500*g* for 1 min and the concentration of released Au NPs was measured by ICP-OES as described above, such that degradation was measured spectroscopically by the cumulative amount of Au released into the media relative to the total amount in as-prepared gelMA-Au NP hydrogels. The remaining hydrogels and/or hydrogel fragments were then weighed on a mass balance, such that degradation was also measured gravimetrically by the total mass change relative to the initial hydrogel mass on day 0. Next, 50 μL DPBS was added to the tube and the hydrogel was imaged by micro-CT, as described above, such that degradation was also measured radiographically by the difference in segmented scaffold volume relative to the initial scaffold volume. Finally, additional fresh DPBS and/or enzyme solution was added back to the tube for continued hydrolytic or enzymatic degradation. Five sample replicates were measured longitudinally for each group.

Enzymatic degradation kinetics were modeled by non-linear lest squares regression using a four-parameter logistic model (GraphPad Prism 9) forcing the *y*-axis from 0 to 100% as,

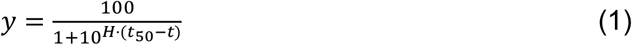

where *y* is the scaffold degradation (%), *t* is time (days), *t*_50_ is the degradation half-life (days), and *H* is the Hill slope or steepness of the kinetic curves which can be used to describe the hydrogel degradation rate. The degradation half-life, Hill slope, and 95% confidence interval were determined from the model. The correlation coefficient (*R*^2^) was greater than 0.9 for each group.

Hydrolytic degradation kinetics measured by micro-CT and ICP-OES were modeled by linear least squares regression (JMP 16). Hydrolytic degradation kinetics measured by gravimetrical analysis were modeled by non-linear least squares regression (GraphPad Prism 9) using the four-parameter logistic model (Eq. 1) with a log-transform of time.

### 2.7. 3D bioprinting of gelMA and gelMA-Au NP hydrogels

Bioinks were prepared with gelMA and one-step photopolymerized gelMA-Au NP prepolymer solutions comprising 20% w/v gelMA, 10 mM AuMA NPs, and 0.5% w/v LAP photoinitiator using methods described above.

#### 2.7.1. Air extrusion bioprinting

Cubic (5 × 5 × 2.5 mm^3^) and two-layer lattice scaffolds with a crosshatch pattern (1 mm^2^ area, 0.3 mm line width, 9 mm strand segment length, and 60° interstrand angle) were printed using a microextrusion bioprinter (BioAssemblyBot, Advanced Solutions Life Sciences). GelMA or gelMA-Au NP inks were loaded in the printing syringe with a 27-gauge tapered needle and printed at room temperature, 6.5 mm/s printing speed, and 0.25 mm line height. Once printed, constructs were immediately photocrosslinked under UV light at 30 mW/cm^2^ for 4 min and subsequently washed with PBS before further evaluation.

#### 2.7.2. Embedded extrusion bioprinting

A 10-layer lattice scaffold (10 × 10 × 2 mm^3^) and hollow cylinder scaffold (6.4 mm outer diameter, 4 mm inner diameter, 2.5 mm in height) were designed and prepared by embedded bioprinting using a microextrusion bioprinter (BioX, CELLINK).^1^ The embedding (support) bath was prepared using 0.4% w/v Carbopol (ETD 2020, Lubrizol) dissolved in DI water. GelMA-Au NP inks were loaded in the printing syringe with a 27-gauge steel needle with a length of 0.5 in and printed at room temperature, 30 kPa printing pressure, and 10 mm/s printing speed. Once printed, constructs were immediately photocrosslinked under UV light at 30 mW/cm^2^ for 3 min and subsequently washed by PBS for 10 min before further evaluation.

#### 2.7.3. Digital light processing (DLP) bioprinting

Three distinct geometric models were created using Fusion360 (Autodesk) and Blender 2.91 (Blender Foundation). These included a cylindrical tube (3 mm outer diameter, 1.5 mm inner diameter), a cube (5 × 5 × 5 mm^3^), and a more complex 3D model of the University of Notre Dame (ND) logo. STL files for each model were processed printed using a DLP bioprinter (Lumen X, CELLINK) with a 0.1 mm layer height, 67% projector power at 10 s/layer (~25 mW/cm^2^), and 2X burn-in on the initial layers.

### 2.8. Characterization of bioprinted gelMA and gelMA-Au NP hydrogel constructs

#### 2.8.1. Printing fidelity

Macroscopic and microscopic (strand-level) dimensional fidelity was assessed on cylindrical tubes and two-layer lattice scaffolds prepared by DLP bioprinting and air extrusion bioprinting, respectively. Construct dimensions measured on digitized optical microscopy images via ImageJ were compared with reference values in the computer-aided design (CAD) model as a ratio, such that a ratio of 1 indicated no deviation. Macroscopic measurements of the cylindrical tube wall thickness ratio, outer diameter ratio, and inner diameter ratio were taken from nine sample replicates. Microscopic measurements of the lattice scaffold strand angle ratio, interstrand area ratio, strand diameter ratio, and strand uniformity ratio^1^ were taken from five sample replicates.

#### 2.8.2. Mechanical properties

Macroscopic and microscopic (strand-level) mechanical properties of cubic gelMA and gelMA-Au NP scaffolds (5 × 5 × 2.5 mm^3^) prepared by air-extrusion bioprinting were measured by unconfined uniaxial compression and microindentation, respectively. Scaffolds were loaded in unconfined uniaxial compression to 50% strain at a displacement rate of 0.02 mm/s using a micromechanical test instrument (Mach-1, Biomomentum). The compressive modulus was calculated as the slope of the stress–strain curve over 10-20% strain using linear least squares regression. For microindentation, scaffolds were loaded on the same instrument with a 500 μm diameter probe to a depth of 100 μm at a displacement rate of 2 μm/s. The reduced elastic modulus was calculated by the Oliver-Pharr method^49^ using the unloading stiffness measured by linear lest squares regression from the initial 10-20% of the unloading force-displacement curves. For each test, three gelMA sample replicates and four gelMA-Au NP sample replicates were measured for each group.

#### 2.8.3. Cell viability in 2D and 3D culture

Human umbilical vein endothelial cells (HUVECs) were cultured in T75 flasks and maintained in complete HUVEC media (VascuLife VEGF Endothelial Medium Complete Kit) supplemented with 1% penicillin/streptomycin (Gibco). HUVEC media was changed every 3 days until cells reached 90% confluency. Cells were then passaged, evaluated, and used for cell culture experiments. For 2D cell culture, a 250 μL prepolymer solution was pipetted into 24-well plates and photocrosslinked under UV light at 10 mW/cm^2^ for 1 min. HUVECs were seeded onto the surface of gelMA-Au NP substrates at 1 × 10^4^ cells/well. After 1 h incubation, 1 mL of HUVEC media was added to each well for cell culture and changed every two days. For 3D cell culture, HUVECs were mixed in gelMA-Au NP inks at a density of 8 × 10^6^ cells/mL. Lattice scaffolds (10 × 10 × 0.2 mm^3^) were printed in an ultra-low attachment 6-well plate by embedded bioprinting using methods described above. Printed scaffolds were photocrosslinked under UV light at 10 mW/cm^2^ for 1 min and immediately cultured in HUVEC media. Cell viability was measured by live/dead assay on days 1 and 7 post-seeding or post-printing. Fluorescent dyes, including 1 μg/mL calcein-AM and 20 μg/mL propidium iodide (Biotium) were added to the culture medium to selectively stain live (green) and dead (red) cells, respectively. After 20 min incubation, substrates or scaffolds were rinsed with fresh culture media and imaged by fluorescence microscopy (DFC3000 G, Leica). Three randomly-selected image fields were evaluated using ImageJ for each of three sample replicates at each time point.

### 2.9. Preparation and characterization of HAMA and HAMA-Au NP hydrogels

HAMA and HAMA-Au NP hydrogels were prepared with AuMA NPs by one-step photocrosslinking to demonstrate the use of AuMA NPs in other photocrosslinkable hydrogels. HAMA with a DoF of 20-50% and molecular weight of 50,000-70,000 HAMA (Sigma-Aldrich) was dissolved in DPBS at 80°C to obtain a 10% w/v HAMA prepolymer solution. HAMA and HAMA-Au NP prepolymer solutions comprising 5% w/v HAMA, 10 mM AuMA NPs, and 0.5% w/v LAP photoinitiator were prepared by mixing the HAMA prepolymer solution with AuMA NPs, vortexing for 2 min, adding LAP, and incubating for 24 h at 4°C. Cylindrical hydrogels were prepared by loading the prepolymer solutions into molds (4.78 mm inner diameter, 3 mm height) and photocrosslinking under UV irradiation (320-390 nm) at 30 mW/cm^2^ for 4 min.

The effective swelling ratio and compressive mechanical properties of HAMA and HAMA-Au NP hydrogels were measured using methods described above. At least four and at most seven sample replicates were tested for each measurement and group.

*In vitro* hydrolytic degradation of HAMA and HAMA-Au NP hydrogels were evaluated longitudinally by micro-CT, ICP-OES, and gravimetric analysis using methods described above, except that micro-CT and gravimetric degradation measurements were normalized by the day 1 time point instead of day 0 because equilibrium swelling was not reached until day 1. Hydrolytic degradation kinetics measured by micro-CT, ICP-OES and gravimetric analysis were modeled by linear least squares regression (JMP 16). At least five and at most seven sample replicates were measured longitudinally for each group.

### 2.10. Statistical Analysis

All quantitative measurements were reported as the mean (± standard deviation) of at least three replicates. Tukey outlier box plots were used to screen for possible outliers defined as data points located outside the interquartile range by more than 1.5-times the interquartile range; one data point was removed for the measured swelling ratio of gelMA hydrogels. The effect of the Au NP concentration on the gelMA-Au NP hydrogel X-ray attenuation was examined using linear least squares regression (JMP 16, SAS Institute). The effect of shear rate on the apparent viscosity of gelMA and gelMA-Au NP prepolymer solutions was also examined using linear least squares regression on a log transform of the data. Differences in the degradation kinetics of hydrogels measured by micro-CT, ICP-OES, and gravimetrical analysis were examined by multivariate linear correlation using Pearson’s correlation coefficient (JMP 16). The effects of synthetic methods and/or hydrogel composition on the rheological properties, swelling ratio, mechanical properties, degradation kinetics, printing fidelity, and cell viability were examined by one-way analysis of variance (ANOVA) and analysis of covariance (ANCOVA). *Post hoc* comparisons were performed using Tukey’s HSD test (JMP 16). Differences in measured printing fidelity ratios were also examined using an exact *t*-test with a hypothesized mean of 1. The level of significance for all tests was set at *p* < 0.05.

## 3. Results and Discussion

### 3.1. Synthesis and characterization of AuMA NPs

AuCOOH NPs were first prepared by the citrate reduction method followed by surface functionalization with MSA.^29,44^ AuMA NPs were then prepared by covalently-linking AuCOOH NPs with AEMA using EDC/NHS chemistry (Fig. 2a). The molar ratio of Au:EDC:NHS:AEMA was set at 1:15:6:6 to maximize the conjugation efficacy of AuCOOH NPs and AEMA, while maintaining solubility and colloidal stability of AuMA NPs in aqueous solution for at least two weeks after synthesis. A higher degree of methacrylation (e.g., Au:EDC:NHS:AEMA = 1:75:30:30) was also investigated, but resulted in hydrophobic NPs which settled out of solution within 2 days after synthesis.

As-prepared AuMA NPs remained spherical and monodispersed with a mean (± standard deviation) diameter of 12.2 (1.2) nm, similar to the initial citrate-stabilized Au NPs and AuCOOH NPs,^29,44^ as measured by TEM (Fig. 2b). Aqueous colloidal stability of AuMA NPs was verified by DLS and zeta potential compared with AuCOOH NPs. The mean (± standard deviation) hydrodynamic diameter of AuMA and AuCOOH NPs was 91.4 (8.0) and 20.3 (1.4) nm (Fig. 2c), and the mean (± standard deviation) zeta potential was −30.9 (1.1) and −13.0 (2.1) mV (Fig. 2d), respectively. The greater hydrodynamic diameter after methacrylation is explained by the association of hydrophobic ligands in aqueous solution, resulting in the formation of dispersible nanoclusters observed in TEM (Fig. 2b). The increased magnitude of negative zeta potential with increased hydrodynamic diameter was previously observed for other functionalized Au NPs.^50^

Methacrylate surface modification was verified by FTIR and NMR spectroscopy of as-prepared AuMA NPs compared with AEMA molecules. FTIR spectra exhibited characteristic C=C (1635 cm^−1^) and =CH (854 cm^−1^) peaks (Fig. 2e), which suggested successful conjugation of AEMA to AuCOOH NPs. ^1^H NMR spectra also confirmed methacrylate surface modification via characteristic peaks at approximately 4.2, 3.7, and 1.9 ppm (Fig. 2f). Characteristic peaks for vinylic protons at approximately 5.7 and 6.1 ppm were not observed for AuMA NPs, but the disappearance and broadening of peaks is commonly observed for small molecules attached to NP surfaces.^51^

### 3.2. Preparation of gelMA-Au NP hydrogels

GelMA-Au NP hydrogels were prepared by one-step photopolymerization (1-step gelMA-Au) with AuMA NPs and compared with gelMA alone, gelMA with physically-entrapped Au NPs (gelMA+Au), and gelMA chemically-coupled with Au NPs using EDC/NHS chemistry prior to photocrosslinking (2-step gelMA-Au) (Fig. 1). AuMA NPs enabled the preparation of chemically-coupled gelMA-Au NP hydrogels in one-step by photopolymerization after simply mixing AuMA NPs with gelMA prepolymer solutions, similar to physical mixing (gelMA+Au). In contrast, chemical-coupling of AuCOOH NPs to gelMA using EDC/NHS chemistry required two separate steps for chemical-coupling and photocrosslinking (2-step gelMA-Au).

#### 3.2.1. Rheology of prepolymer solutions

Prepolymer solutions for 1-step gelMA-Au hydrogels exhibited the most similar rheological properties compared with gelMA alone (Fig. S1). Both 1-step gelMA-Au and gelMA+Au prepolymer solutions did not alter the sol-gel transition temperature of gelMA (Fig. S1a,b). In contrast, 2-step gelMA-Au prepolymer solutions lost a thermal sol-gel transition when efficient chemical-coupling of Au-COOH NPs to gelMA macromolecules was assured by using an excess of EDC/NHS. Two-step gelMA-Au prepolymer solutions prepared with a Au:EDC:NHS molar ratio of 1:160:64 did not exhibit a thermal sol-gel transition when heated as high as 80°C, most likely due EDC/NHS-mediated crosslinking of amine and carboxylate ligands on gelMA macromolecules. Two-step gelMA-Au prepolymer solutions prepared with a Au:EDC:NHS molar ratio of 1:1:1 were able to achieve a sol-gel transition temperature comparable to gelMA alone (Fig. S1a,b), but only with compromised efficiency for chemical-coupling AuCOOH NPs to gelMA and disruption of the photocrosslinked hydrogel network, as shown below.

GelMA and gelMA-Au NP prepolymer solutions all exhibited shear thinning with increased shear rate (*p* < 0.0.001, ANCOVA), as expected (Fig. S1c). One-step gelMA-Au prepolymer solutions exhibited a lower apparent viscosity compared with gelMA overall (*p* < 0.05, Tukey), but meaningful differences were primarily at low shear rates (< 10 s^−1^). At higher shear rates (> 10 s^−1^) relevant to 3D printing processes, gelMA and 1-step gelMA-Au prepolymer solutions exhibited similar apparent viscosity (Fig. S1c). In contrast, gelMA+Au and 2-step gelMA-Au prepolymer solutions exhibited lower apparent viscosity compared with both gelMA and 1-step gelMA-Au prepolymer solutions overall (*p* < 0.05, Tukey), and especially at higher shear rates.

#### 3.2.2. Optimization of photocrosslinking

The concentration of Au NPs in gelMA-Au NP hydrogels was recognized to exhibit competing effects on X-ray contrast and photocrosslinking. Greater concentrations of Au NPs enable greater absorption of X-rays,^31^ which is beneficial for radiographic contrast, but also increase the absorption and scattering of UV and visible light,^52^ which inhibits photocrosslinking (Table S1). Therefore, photocrosslinking studies were performed to optimize key parameters – including the Au NP concentration; photoinitiator type, concentration, and incubation time; gelMA concentration; and UV light intensity and irradiation time – for achieving fully crosslinked hydrogels with sufficient radiographic contrast, while minimizing the UV irradiation intensity and time to avoid damage to encapsulated cells and/or biomolecules.

GelMA+Au NP hydrogels prepared with 10% w/v gelMA, 0.5% w/v Irgacure photoinitiator, and 0-22 mM Au NPs initially revealed that hydrogels containing up to 7 mM Au NPs were fully crosslinked and hydrogels containing at least 15 mM Au NPs exhibited little or no crosslinking under UV irradiation at 7 and 15 mW/cm^2^ for 6 min (Table S1). Therefore, a subsequent study systematically investigated increased photoinitiator concentration (1.0% w/v Irgacure) and UV light intensity (30 mW/cm^2^, 4 min), but gelMA+Au NP hydrogels containing more than 7 mM Au NPs were still not fully crosslinked (Table S2). These results, and the inability to improve photocrosslinking with increased photoinitiator concentration and UV light intensity, suggested that the effective UV intensity for activating the photoinitator was reduced by absorption and scattering from Au NPs.

One-step gelMA-Au NP hydrogels prepared under conditions previously shown to result in little or no crosslinking in gelMA+Au NP hydrogels (Table S2) also exhibited little or no photocrosslinking using 10% w/v gelMA, but exhibited improved crosslinking using 20% w/v gelMA, with either 15 or 37 mM Au NPs (Table S3). The increased gelMA concentration increased the number of potential crosslinking sites and their proximity with the photoinitiator without a loss in transparency for UV light penetration. Finally, lithium phenyl-2,4,6-trimethylbenzoylphosphinate (LAP) was investigated as an alternative photoinitiator known to exhibit more rapid photopolymerization kinetics and higher water solubility compared to Irgacure 2959.^47,53^ LAP improved photocrosslinking of 1-step gelMA-Au hydrogels prepared with 20% w/v gelMA and up to 37 mM Au NPs compared to Irgacure, especially after incubating prepolymer solutions with the photoinitiator for at least 24 h prior to photocrosslinking (Table S4). This result suggests that in addition to absorbing and scattering of UV light AuMA NPs may have inhibited photocrosslinking due to increasing the hydrophobicity prepolymer solutions. Importantly, 1-step gelMA-Au hydrogels comprising up to 10 and 37 mM AuMA NPs were fully crosslinked after incubating prepolymer solutions with 0.5% w/v LAP for 24 h and 7 d, respectively, and photocrosslinking under UV irradiation at 30 mW/cm^2^ for 4 min (Table S4).

In summary, challenges to achieving fully crosslinked gelMA-Au NP hydrogels were circumvented by increasing the gelMA concentration, adopting LAP as the photoinitiator, and incubating prepolymer solutions with LAP for at least 24 h prior to photocrosslinking. GelMA and gelMA-Au NP hydrogels described hereafter were prepared with 20% w/v gelMA and up to 37 mM Au NPs by incubating prepolymer solutions with 0.5% w/v LAP photoinitiator 24 h at 4°C and photocrosslinking under UV irradiation at 30 mW/cm^2^ for 4 min.

### 3.3. AuMA NPs provide tunable X-ray contrast in photopolymerized gelMA hydrogels

GelMA-Au NP hydrogels were prepared by the one-step photocrosslinking strategy with varying concentrations of AuMA NPs to determine a suitable Au NP concentration for longitudinal monitoring of hydrogels by micro-CT. The X-ray attenuation of as-prepared 1-step gelMA-Au hydrogels increased linearly with increasing Au NP concentration(*p* < 0.001) and was strongly correlated (*R*^2^ = 0.99), as expected (Fig. 3a). This result confirmed that AuMA NPs can provide tunable X-ray contrast in gelMA hydrogels to enable quantitative imaging by micro-CT. GelMA-Au NP hydrogels containing at least 5 mM Au NPs exhibited sufficient X-ray attenuation for contrast enhancement (△HU ≥ 30^54^) versus soft tissue (Fig. 3a), as represented by PBS (38.3 HU) and myocardium tissue (−9.3 HU). Grayscale micro-CT images of hydrogels containing at least 5 mM Au NPs exhibited visibly greater X-ray attenuation compared with PBS (Fig. 3a). A Au NP concentration of 10 mM was therefore chosen to provide sufficient X-ray contrast versus soft tissue for monitoring degradation.

**Fig. 3.**
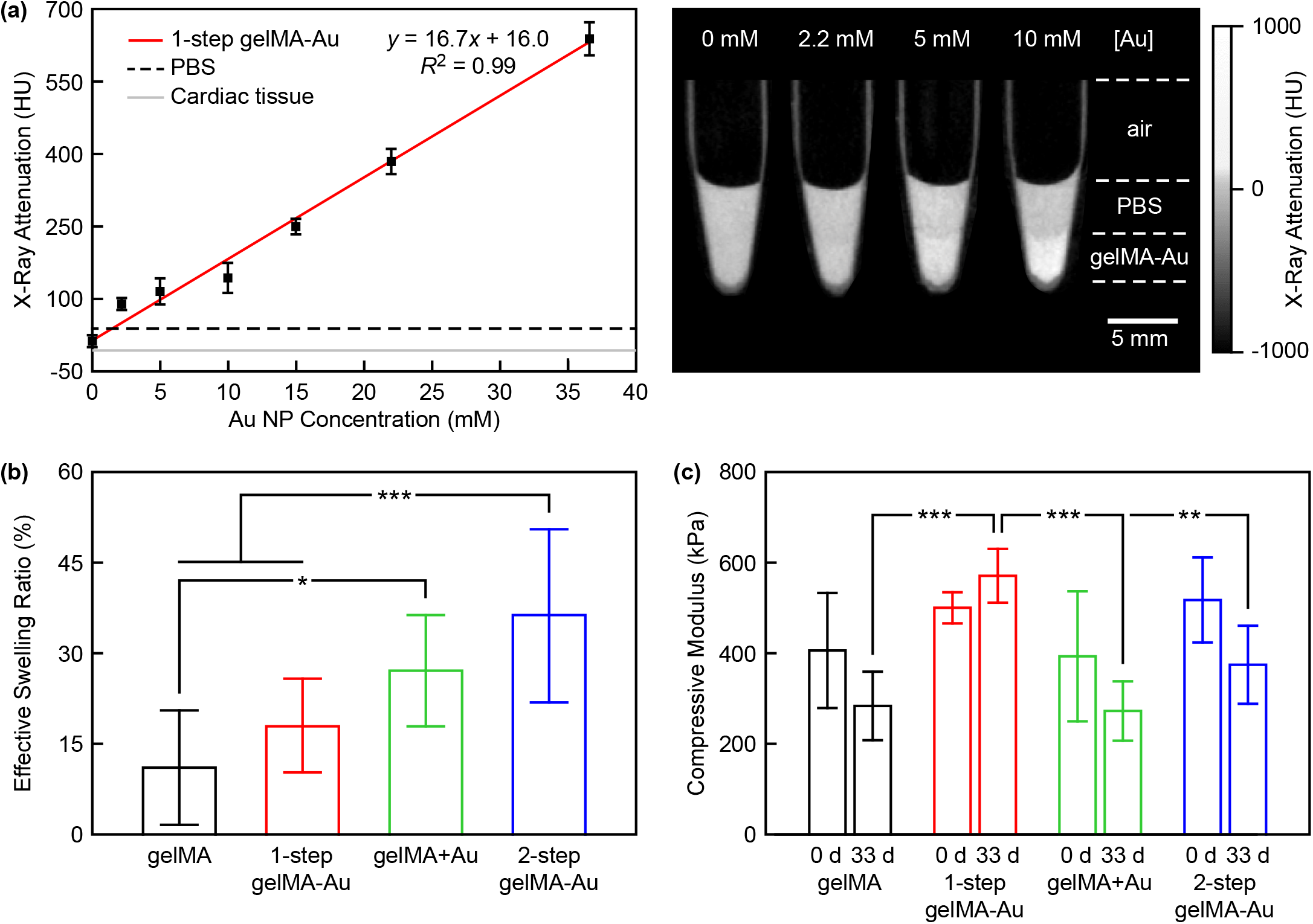
Characterization of gelMA-Au NP hydrogels. (a) The X-ray attenuation of gelMA-Au NP hydrogels prepared with varying concentration of AuMA NPs by one-step photopolymerization (1-step gelMA-Au) compared with soft tissue, as represented by PBS (38.3 HU) and rat myocardial tissue (−9.3 HU). The measured X-ray attenuation increased linearly with increased Au NP concentration (*p* < 0.001) and was strongly correlated (*R*^2^ = 0.99). Error bars show one standard deviation of the mean (*n* = 3/concentration). Grayscale micro-CT images showed that hydrogels containing at least 5 mM Au NPs exhibited visibly greater X-ray attenuation compared with PBS. (b) The effective swelling ratio of gelMA and gelMA-Au NP hydrogels after reaching equilibrium swelling (*n* = 10/group). The effective swelling ratio of 1-step gelMA-Au hydrogels was not statistically different from gelMA alone (*p* > 0.48, Tukey), but was greater for other gelMA-Au hydrogels compared with gelMA. (c) The compressive modulus of gelMA and gelMA-Au NP hydrogels measured before (0 d) and after 33 days hydrolysis (*n* = 3-5/group/time point). The compressive modulus of as-prepared gelMA and gelMA-Au hydrogels was not different (*p* > 0.41, ANOVA), but was maintained and greater for 1-step gelMA-Au hydrogels, compared to other groups after 33 days hydrolysis. Error bars show one standard deviation of the mean. \**p* < 0.05, \*\**p* < 0.01, \*\*\**p* < 0.005, Tukey.

### 3.4. AuMA NPs do not disrupt the gelMA hydrogel network, mechanical properties, and hydrolytic stability

The effective swelling ratio of gelMA-Au NP hydrogels was measured and compared to gelMA alone to evaluate the hydrogel crosslinking density. The effective swelling ratio of 1-step gelMA-Au hydrogels was not statistically different (*p* > 0.48, Tukey) from gelMA alone (Fig. 3b). In contrast, physically-mixed gelMA+Au hydrogels and 2-step gelMA-Au hydrogels exhibited a greater swelling ratio compared to gelMA alone (*p* < 0.05, Tukey). These results suggest that 1-step gelMA-Au hydrogels prepared with AuMA NPs did not disrupt the hydrogel network or crosslinking density compared with gelMA alone. AuMA NPs provided numerous ligands for photocrosslinking into the gelMA hydrogel network (Fig. 1a). In contrast, physically-mixed gelMA+Au hydrogels and 2-step gelMA-Au hydrogels disrupted the hydrogel network and/or reduced the crosslinking density compared with gelMA alone. In physically-mixed gelMA+Au hydrogels, AuCOOH NPs were unable to participate in photocrosslinking and thus disrupted the gelMA hydrogel network (Fig. 1b). In 2-step gelMA-Au hydrogels, chemical-coupling of AuCOOH NPs to gelMA macromolecules prior to photocrosslinking not only disrupted the gelMA hydrogel network but also likely decreased the crosslinking density of gelMA-Au macromolecules compared with gelMA macromolecules (Fig. 1c).

The compressive modulus of gelMA-Au hydrogels was measured and compared to gelMA alone before (day 0) and after 33 days of hydrolysis in DPBS to evaluate the hydrogel crosslinking density and hydrolytic stability. The compressive modulus of gelMA and gelMA-Au hydrogels was not statistically different (*p* > 0.41, ANOVA) at day 0 (Fig. 3c). However, following 33 days of hydrolysis, the compressive modulus of 1-step gelMA-Au hydrogels was maintained at a similar level as day 0, but greater (*p* < 0.01, Tukey) than that for gelMA, gelMA+Au, and 2-step gelMA-Au hydrogels (Fig. 3c). Taken together, these results suggest that 1-step gelMA-Au hydrogels prepared with AuMA NPs did not disrupt the initial mechanical properties of gelMA hydrogels and were more resistant to hydrolytic degradation. Additionally, physically-mixed gelMA+Au hydrogels and 2-step gelMA-Au hydrogels exhibited a greater effective swelling ratio (Fig. 3b) but unchanged compressive modulus (Fig. 3c) compared with gelMA alone. These results suggest that the Au NPs provided mechanical reinforcement that compensated for a disrupted hydrogel network and/or reduced crosslinking density. Supporting this observation, physically-entrapped gelMA+Au hydrogels were previously shown to exhibit a decreased swelling ratio and increased compressive modulus with increased concentration of Au nanorods.^37^

The hydrolytic stability of gelMA-Au NP hydrogels was also monitored longitudinally by contrast-enhanced micro-CT, ICP-OES, and gravimetric analysis. All gelMA and gelMA-Au NP hydrogels were stable against hydrolysis for at least one month (Fig. S2). Differences in the cumulative degradation between gelMA and gelMA-Au NP hydrogels after 30 days hydrolysis were not statistically significant (*p* > 0.62, ANOVA). However, the cumulative degradation measured by gravimetric analysis was greater than that measured by micro-CT and ICP-OES. For example, gelMA-Au hydrogels prepared by 1-step photocrosslinking with AuMA NPs exhibited a cumulative degradation of ~5% as measured by micro-CT and ICP-OES, and ~20% as measured by gravimetric analysis, after 30 days hydrolysis (Fig. S2). The greater degradation measured by gravimetric analysis occurred nearly entirely within the first three days and was unchanged thereafter. The initial ~15% loss of mass within the first three days (Fig. S2d) was not accompanied by release of Au NPs (Fig. S2c). Taken together, these results suggest that gelMA and gelMA-Au NP hydrogels exhibited thermally-induced deswelling upon incubation at 37°C^55^ which caused an initial loss of mass as hydrogels re-equilibrated, and not hydrolytic degradation.

### 3.5. AuMA NPs enable longitudinal monitoring of gelMA hydrogels during enzymatic degradation

Enzymatic degradation of gelMA-Au NP hydrogels was monitored longitudinally *in vitro* for more than three weeks by contrast-enhanced micro-CT and compared with concurrent measurements by gravimetric analysis and optical spectroscopy (Fig. 4). Segmented micro-CT image reconstructions showed clear and repeatable changes in the hydrogel volume over time due to enzymatic degradation (Fig. 4a). Importantly, contrast-enhanced micro-CT was able to quantitatively measure the degradation kinetics and differences in the degradation kinetics between 1-step gelMA-Au, physically-mixed gelMA+Au NP, and 2-step gelMA-Au hydrogels (Fig. 4b, Table 1). The degradation kinetics measured by contrast-enhanced micro-CT (Fig. 4b) exhibited close agreement with that measured by ICP-OES (Fig. 4c) and gravimetric analysis (Fig. 4d). Direct comparison of the degradation kinetics of 1-step gelMA-Au hydrogels measured by each technique (Fig. 4e) revealed that the degradation kinetics measured by contrast-enhanced micro-CT were strongly correlated (*r* > 0.96, Pearson) with that measured by ICP-OES and gravimetric analysis. These results demonstrate the feasibility of contrast-enhanced micro-CT for non-invasive monitoring of gelMA-Au NP hydrogel degradation.

**Fig. 4.**
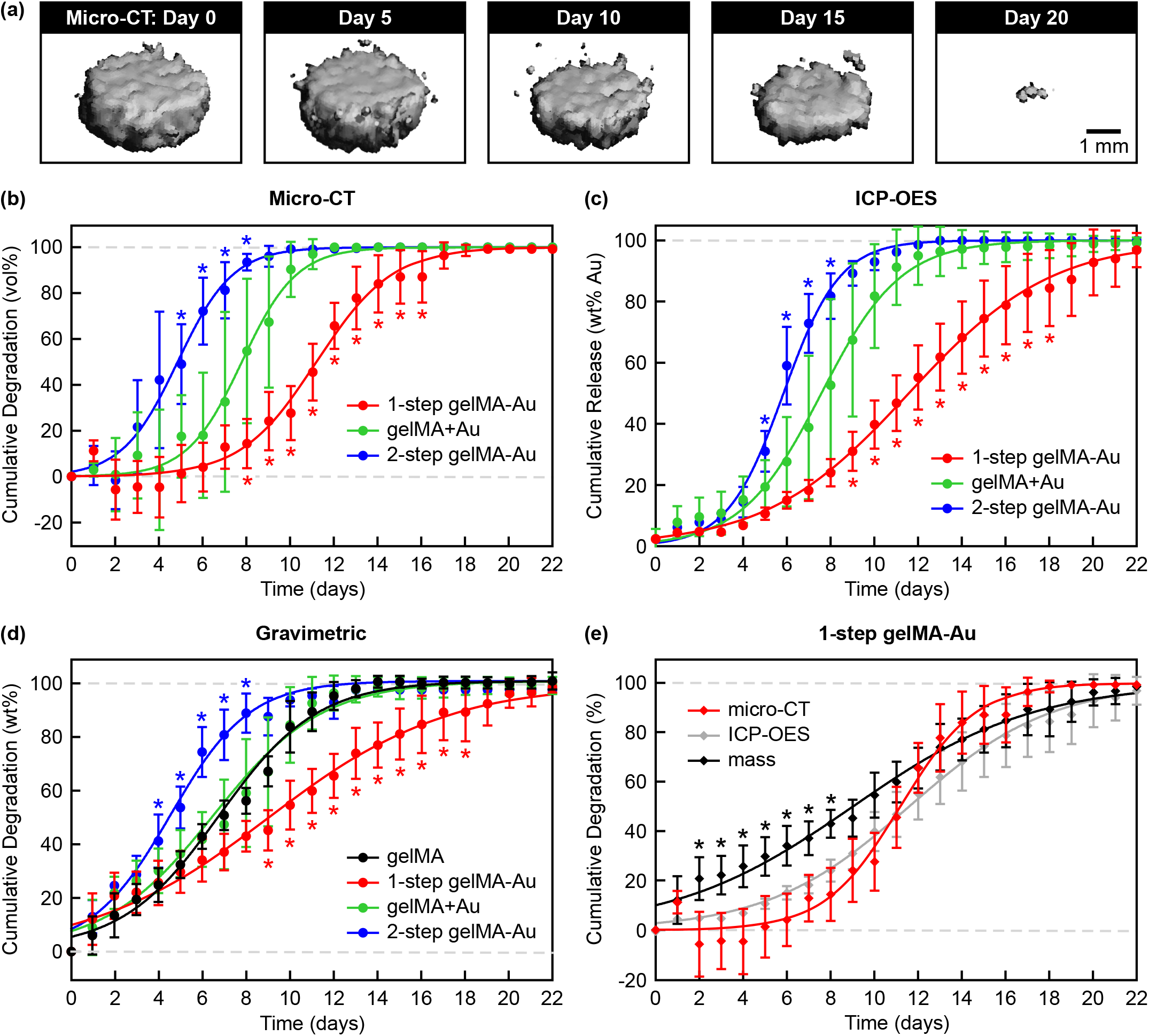
Non-invasive monitoring of gelMA and gelMA-Au NP hydrogels during *in vitro* enzymatic degradation by contrast-enhanced micro-CT. (a) Representative segmented micro-CT image reconstructions for selected time points showing the volume loss of 1-step gelMA-Au hydrogels during enzymatic degradation. Degradation kinetics were measured longitudinally by (b) the cumulative change in segmented hydrogel volume using contrast-enhanced micro-CT, (c) the cumulative release of Au NPs into the media using ICP-OES, and (d) the cumulative hydrogel mass loss using gravimetric analysis. 1-step gelMA-Au hydrogels exhibited slower degradation kinetics, while 2-step gelMA-Au hydrogels exhibited more rapid degradation kinetics, compared with gelMA and gelMA+Au hydrogels. (e) Comparison of the degradation kinetics for 1-step gelMA-Au hydrogels measured longitudinally by micro-CT, ICP-OES, and gravimetric analysis. The degradation kinetics measured by micro-CT were strongly correlated (*r* > 0.96, Pearson) with that measured by ICP-OES and gravimetric analysis, demonstrating the feasibility of contrast-enhanced micro-CT for non-invasive monitoring of gelMA-Au NP hydrogel degradation. Degradation kinetics were modeled by non-linear least squares regression using a four-parameter logistic model (Eq. 1) with fitting parameters reported in Table 1. Error bars show one standard deviation of the mean (*n* = 5/group). Error bars not shown lie within the data point. \**p* < 0.05 vs. gelMA+Au in (b,c), vs. gelMA in (d), vs. micro-CT and ICP-OES in (e) Tukey.

**Table 1.** Non-linear least squares regression of gelMA and gelMA-Au NP hydrogel enzymatic degradation kinetics using a four-parameter logistic model (Eq. 1), where *t*_50_ is the degradation half-life, including the 95% confidence interval (CI), *H* is the Hill slope or steepness of the kinetic curves which reflects the hydrogel degradation rate, and *R*^2^ is the model correlation coefficient.

| Group | Test Method | $t_{50}$ (days) | $t_{50}$, 95% CI | $H$ | $R^2$ |
| --- | --- | --- | --- | --- | --- |
| gelMA | gravimetric | 6.8 | 6.6 - 7.0 | 0.19 | 0.98 |
| 1-step gelMA-Au | micro-CT | 11.2 | 10.9 - 11.5 | 0.25 | 0.95 |
|  | ICP-OES | 11.6 | 11.3 - 11.9 | 0.13 | 0.95 |
|  | gravimetric | 9.0 | 8.7 - 9.4 | 0.11 | 0.94 |
| gelMA+Au | micro-CT | 7.8 | 7.4 - 8.1 | 0.35 | 0.89 |
|  | ICP-OES | 7.6 | 7.3 - 7.9 | 0.24 | 0.92 |
|  | gravimetric | 6.5 | 6.2 - 6.8 | 0.18 | 0.93 |
| 2-step gelMA-Au | micro-CT | 4.8 | 4.6 - 5.1 | 0.34 | 0.94 |
|  | ICP-OES | 5.9 | 5.8 - 6.0 | 0.34 | 0.99 |
|  | gravimetric | 4.5 | 4.4 - 4.7 | 0.24 | 0.97 |

1-step gelMA-Au hydrogels exhibited slower degradation kinetics, while 2-step gelMA-Au hydrogels exhibited more rapid degradation kinetics, compared with gelMA and gelMA+Au hydrogels (Fig. 4, Table 1). The slower degradation kinetics exhibited by 1-step gelMA-Au hydrogels was most likely the result of AuMA NPs participating in photocrosslinking to form additional crosslinks within the hydrogel network (Fig. 1a). A common limitation of hydrogels is rapid degradation and/or burst release of encapsulated drugs.^39,56,57^ Therefore, these results suggest that this limitation can be mitigated by incorporating AuMA NPs in photocrosslinked hydrogels. In contrast, the more rapid degradation kinetics exhibited by 2-step gelMA-Au hydrogels was most likely the result of Au NPs covalently-linked to gelMA molecules prior to photocrosslinking disrupting the hydrogel network and reducing the crosslinking density during photocrosslinking (Fig. 1c), consistent with results for the effective swelling ratio and mechanical properties (Fig. 3b,c).

Gravimetric measurements exhibited more rapid degradation at early time points than concurrent measurements by contrast-enhanced micro-CT and ICP-OES (Fig. 4b-d). For example, the cumulative degradation of 1-step gelMA-Au hydrogels measured by gravimetric analysis was greater than that measured by contrast-enhanced micro-CT and ICP-OES at days 2-8 (*p* < 0.05, Tukey) while differences between micro-CT and ICP-OES were not statistically significant at any time point (Fig. 4e). These differences were reflected in measurements of the degradation half-life (*t*_50_) and rate (*H*) (Table 1). Gravimetric analysis cannot distinguish mass lost due to deswelling versus degradation of the hydrogel network. Therefore, these results suggest that measurements by contrast-enhanced micro-CT and ICP-OES more likely reflected the true degradation kinetics. The close agreement between micro-CT and ICP-OES further suggests that contrast-enhanced micro-CT enabled precise and accurate measurement of hydrogel degradation non-invasively.

### 3.6. 3D bioprinting gelMA bioinks with AuMA NPs to form 1-step gelMA-Au hydrogel constructs

GelMA bioinks supplemented with AuMA NPs were printed by extrusion (Fig. 5a, S3a) and DLP (Fig. 5b, S3a) bioprinting. As noted above, gelMA prepolymer solutions supplemented with AuMA NPs exhibited similar rheological properties as gelMA alone (Fig. S1) and shear-thinning behavior under flow, which is advantageous for high-fidelity extrusion bioprinting.^1,58^

**Fig. 5.**
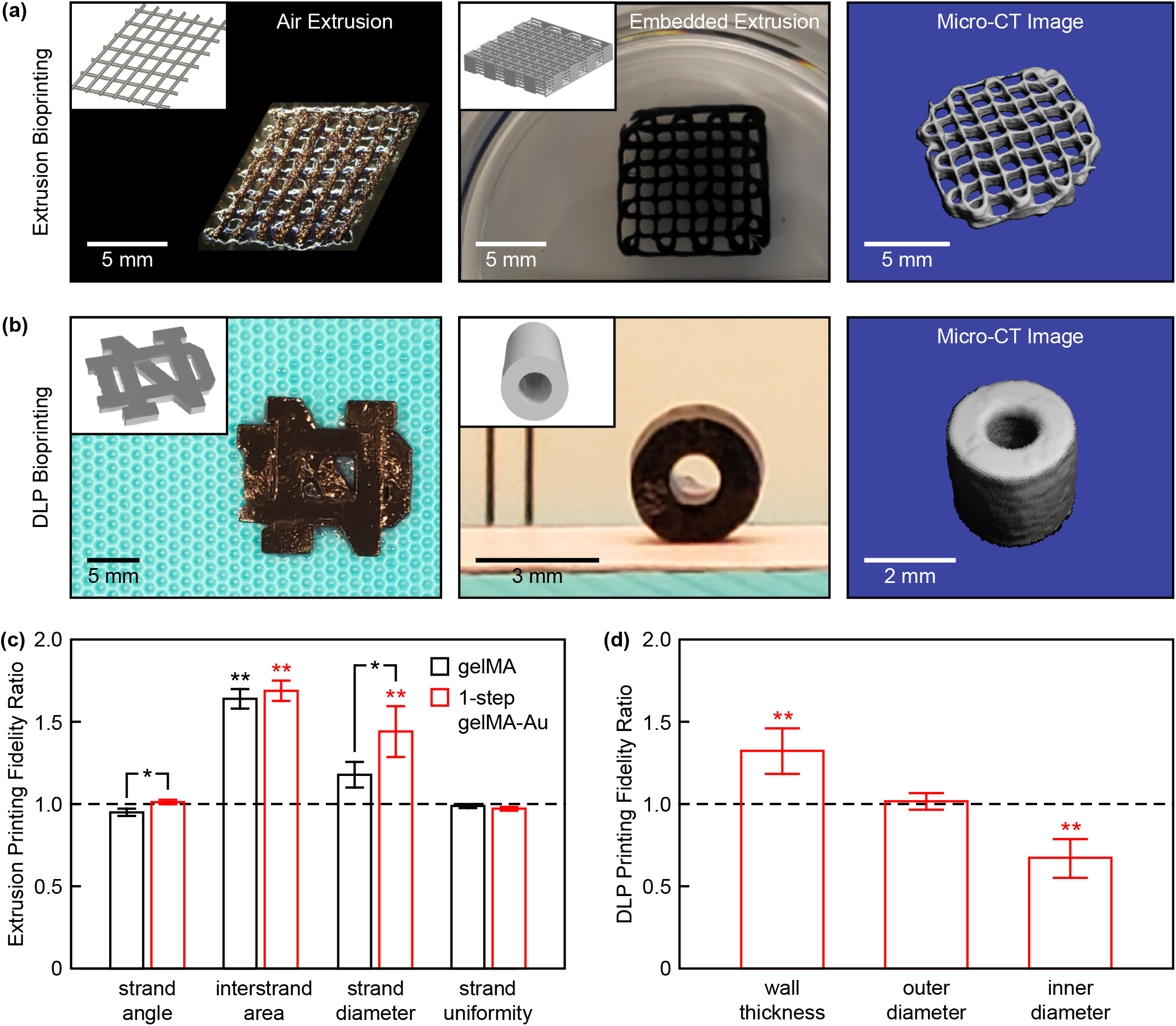
3D bioprinting of 1-step gelMA-Au NP hydrogel constructs. (a) A two-layer lattice scaffold printed by air extrusion and 10-layer lattice scaffold printed by embedded extrusion. (b) The University of Notre Dame logo (left) and a cylindrical tube mimicking a blood vessel printed by DLP bioprinting. Insets show corresponding CAD models. Segmented micro-CT image reconstructions of the 10-layer lattice scaffold and cylindrical tube show feasibility of non-invasive radiograph imaging. (c) Microscale printing fidelity measured for two-layer lattice scaffolds printed by air extrusion with gelMA and 1-step gelMA-Au hydrogels. The printing fidelity of 1-step gelMA-Au hydrogels was comparable to that of gelMA alone. Error bars show one standard deviation of the mean (*n* = 5/group). (d) Macroscale printing fidelity measured for cylindrical tubes printed by DLP bioprinting with 1-step gelMA-Au hydrogels. Error bars show one standard deviation of the mean (*n* = 9/group). In (c) and (d), a ratio of one (dashed line) indicates no deviation from the CAD models. \**p* < 0.05, Tukey. \*\**p* < 0.005 vs. 1, exact *t*-test.

#### 3.6.1. Printing fidelity

Two-layer lattice (Fig. 5a) and cubic (Fig. S3a) scaffolds were printed by air extrusion with adequate reproducibility in correspondence to the CAD models. The microscale printing fidelity of two-layer lattice scaffolds was quantified by the ratio of printed construct dimensions relative to the CAD model. Overall, the printing fidelity of 1-step gelMA-Au hydrogels was comparable to that of gelMA alone with only small differences in the strand angle and diameter (Fig. 5c). Differences in the strand angle and uniformity between printed constructs and the CAD model were not statistically significant for both gelMA and 1-step gelMA-Au hydrogels. Differences in the interstrand area and strand diameter between printed constructs and the CAD model were most likely caused by viscoelastic flow after printing (creep) and die swell during extrusion (stress relaxation), respectively.^1,59^ More complex constructs, including 10-layer lattice (Fig. 5a) and hollow cylinder (Fig. S3a) scaffolds, were also printed by embedded extrusion, which was previously shown to enable improved printing fidelity compared with air extrusion printing.^1^

DLP bioprinting was utilized to print 1-step gelMA-Au hydrogel constructs of varying 3D geometry, including a University of Notre Dame logo (Fig. 5b), cylindrical tubes mimicking blood vessels (Fig. 5b), and a cubic stucture (Fig. S4a). The macroscale printing fidelity of cylindrical tubes was quantified by the ratio of printed construct dimensions relative to the CAD model. Differences in the outer diameter between printed constructs and the CAD model were not statistically significant (Fig. 5d). Differences in the inner diameter and corresponding wall thickness between printed constructs and the CAD model (*p* < 0.005, exact *t*-test) were most likely due to incomplete photocrosslinking caused by the absorption and scattering of light by Au NPs in the thick cross-section. Thus, further optimization of printing and photocrosslinking parameters is required. Nonetheless, for both extrusion and DLP bioprinting, dimensional differences between constructs and CAD models were consistent and can therefore be compensated by scaling dimensions in STL files.^60^ Overall, gelMA bioinks supplemented with AuMA NPs exhibited suitable printability and dimensional fidelity.

#### 3.6.2. Contrast-enhanced micro-CT

Segmented micro-CT image reconstructions of the 10-layer lattice (Fig. 5a) and cylindrical tube (Fig. 5b) scaffolds demonstrated feasibility of non-invasive radiographic imaging of bioprinted constructs. AuMA NPs provided sufficient X-ray contrast for imaging both the macrostructure and microstructural features, whereas constructs printed from gelMA alone could not be imaged by micro-CT.

#### 3.6.3. Mechanical properties

The mechanical integrity of gelMA and gelMA-Au hydrogels printed by air extrusion was evaluated on cubic constructs in unconfined uniaxial compression and microindentation after reaching equilibrium swelling (Fig. S3). Differences in the compressive modulus and indentation modulus between gelMA and 1-step gelMA-Au hydrogels were not statistically significant. Therefore, AuMA NPs did not disrupt the mechanical properties of bioprinted 1-step gelMA-Au hydrogels, consistent with measurements for molded 1-step gelMA-Au hydrogels (Fig. 3c).

### 3.7. Bioprinted 1-step gelMA-Au NP hydrogels support cell viability in 2D and 3D culture

Cell viability in 1-step gelMA-Au hydrogels was assessed by both 2D and 3D culture of HUVECs in bioprinted constructs (Fig. 6). In 2D culture, over 95% of cells seeded onto the surface of 1-step gelMA-Au hydrogel substrates remained viable at days 1 and 7. In 3D culture, over 70% of cells that were encapsulated in the bioink before printing remained viable within 1-step gelMA-Au hydrogels at days 1 and 7. Decreased cell viability in 3D culture is commonly observed in bioprinted constructs due to cellular damage caused by mechanical forces introduced to cells during extrusion and/or UV light exposure during crosslinking.^61,62^ Printing-induced damage to cells can be reduced using a lower printing pressure or speed, larger gauge syringe needle, or lower UV intensity or exposure time. Limited nutrient transport in 3D culture, especially hydrogels prepared with high gelMA concentration, may have also contributed to decreased cell viability.^63^ Importantly, however, there was no loss of viability between days 1 and 7 (*p* > 0.71, ANOVA) in either 2D or 3D culture. Moreover, HUVECs exhibited a more elongated and mature morphology at day 7 compared with day 1 in both 2D and 3D culture. Taken together, these results suggest that 1-step gelMA-Au hydrogels are cytocompatible with endothelial cells.

**Fig. 6.**
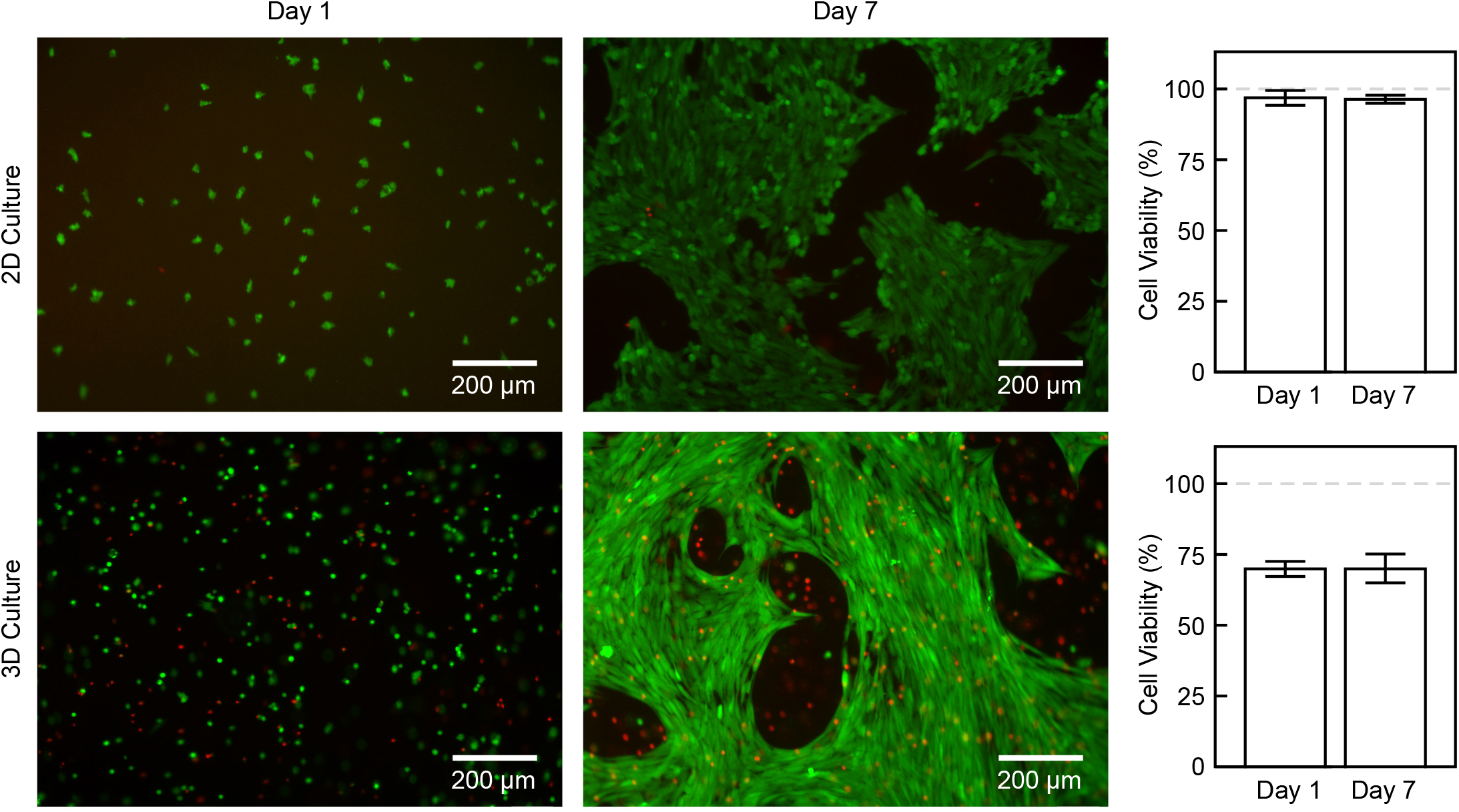
Cell viability in 3D bioprinted 1-step gelMA-Au NP hydrogel constructs. Representative epifluorescence micrographs showing live(green)/dead(red) staining of HUVECs after 1 and 7 days of 2D and 3D culture with 1-step gelMA-Au hydrogels. Quantitative measurements showed more than 95% viable cells in 2D culture and more than 70% viable cells in 3D culture. There was no loss of viability over 7 days culture (*p* > 0.71, ANOVA).

### 3.8. AuMA NPs are applicable to other photopolymerized hydrogels

HAMA-Au hydrogels were also prepared with AuMA NPs by one-step photocrosslinking to demonstrate the use of AuMA NPs in other photocrosslinkable hydrogels. The effective swelling ratio of as-prepared HAMA-Au hydrogels was increased compared with HAMA alone (*p* < 0.05, Tukey) (Fig. S5a). This result suggests that, unlike in gelMA hydrogels, AuMA NPs decreased the crosslinking density of HAMA hydrogels, most likely due to the lower methacrylation degree of HAMA (20-50%) versus gelMA (80%) in this study. The compressive modulus of HAMA and HAMA-Au hydrogels was not significantly different (*p* > 0.75, ANOVA) (Fig. S5b). This result suggests that the Au NPs provided mechanical reinforcement that compensated for the reduced crosslinking density, which is consistent with the results for gelMA-Au hydrogels (Fig. 3b,c). Both HAMA and 1-step HAMA-Au hydrogels were stable against hydrolysis for at least four weeks (Fig. S4c-f). Differences in the cumulative degradation between HAMA and 1-step HAMA-Au hydrogels were not statistically significant (Fig. S4f). Differences in the cumulative degradation of 1-step HAMA-Au hydrogels measured by contrast-enhanced micro-CT, ICP-OES, and gravimetric analysis were also not statistically significant. Taken together, these results suggest that HAMA and other methacrylate-modified hydrogels are also readily prepared with AuMA NPs by one-step photocrosslinking to enable non-invasive monitoring by contrast-enhanced micro-CT.

## 4. Conclusions

Methacrylate-modified gold nanoparticles (AuMA NPs) were prepared and covalently-linked to methacrylate-modified hydrogels (gelMA and HAMA) in one-step during photocrosslinking. Photopolymerized hydrogels exhibited a linear increase in X-ray attenuation with increased Au NP concentration to enable quantitative imaging by contrast-enhanced micro-CT. The hydrolytic and enzymatic degradation kinetics of gelMA-Au NP hydrogels were longitudinally monitored by micro-CT for up to one month *in vitro*, and were consistent with concurrent measurements by gravimetric analysis and optical spectroscopy. Importantly, AuMA NPs provided little or no disruption to the hydrogel network, rheology, mechanical properties, and hydrolytic stability compared with gelMA alone. GelMA-Au NP hydrogels were able to be printed into well-defined 3D architectures supporting endothelial cell viability. Thus, AuMA NPs enabled the preparation of photopolymerized hydrogels and bioprinted scaffolds with tunable X-ray contrast for noninvasive, longitudinal monitoring of degradation and NP release by micro-CT.

## Author contributions

L.L., C.J.G., V.S and R.K.R. designed experiments, analyzed data, and wrote the manuscript. L.L. and C.J.G. were primarily responsible for performing all experiments. T.A.F., C.J.E., L.N., M.L.T., A.T., and G.K. provided assistance in performing experiments and editing the manuscript.

## Acknowledgments

L.L. was supported by a Materials Science and Engineering Program Doctoral Fellowship from the University of Notre Dame. C.J.G was supported by a National Science Foundation (NSF) Graduate Research Fellowship (GE-1650044) and the American Heart Association (AHA) Predoctoral Fellowship (20PRE35080132). C.J.E. was supported by the Martell Family Ph.D. Fellowship at the University of Notre Dame. The authors acknowledge additional support from the Kelly Cares Foundation, National Institutes of Health (R00HL127295), Pediatric Research Alliance, and Emory University Dean’s Imagine, Innovate and Impact (I3) Research Award. The authors acknowledge the Center for Environmental Science and Technology (CEST) for use of ICP-OES and FTIR, the Materials Characterization Facility (MCF) for rheometry, and Dr. Pinar Zorlutuna for providing the myocardial tissue sample, all from the University of Notre Dame.

## Appendix A. Supplementary data

Supplementary data to this article can be found online.

## Data Availability

The raw data required to reproduce these findings are available to download from doi:10.17632/gnv5k2hb8t.1

## References

1. L. Ning, R. Mehta, C. Cao, A. Theus, M. Tomov, N. Zhu, E.R. Weeks, H. Bauser-Heaton, V. Serpooshan, Embedded 3D bioprinting of gelatin methacryloyl-based constructs with highly tunable structural fidelity, ACS Appl. Mater. Interfaces 12 (2020) 44563–44577. https://doi.org/10.1021/acsami.0c15078.

2. B. Kong, Y. Chen, R. Liu, X. Liu, C. Liu, Z. Shao, L. Xiong, X. Liu, W. Sun, S. Mi, Fiber reinforced GelMA hydrogel to induce the regeneration of corneal stroma, Nat. Commun. 11 (2020) 1435. https://doi.org/10.1038/s41467-020-14887-9.

3. W. Schuurman, P.A. Levett, M.W. Pot, P.R. van Weeren, W.J.A. Dhert, D.W. Hutmacher, F.P.W. Melchels, T.J. Klein, J. Malda, Gelatin-methacrylamide hydrogels as potential biomaterials for fabrication of tissue-engineered cartilage constructs, Macromol. Biosci. 13 (2013) 551–561. https://doi.org/10.1002/mabi.201200471.

4. J.A. Burdick, G.D. Prestwich, Hyaluronic acid hydrogels for biomedical applications, Adv. Mater. 23 (2011) H41–H56. https://doi.org/10.1002/adma.201003963.

5. J.L. Ifkovits, E. Tous, M. Minakawa, M. Morita, J.D. Robb, K.J. Koomalsingh, J.H. Gorman, R.C. Gorman, J.A. Burdick, Injectable hydrogel properties influence infarct expansion and extent of postinfarction left ventricular remodeling in an ovine model, Proc. Natl. Acad. Sci. 107 (2010) 11507–11512. https://doi.org/10.1073/pnas.1004097107.

6. J. Hjortnaes, G. Camci-Unal, J.D. Hutcheson, S.M. Jung, F.J. Schoen, J. Kluin, E. Aikawa, A. Khademhosseini, Directing valvular interstitial cell myofibroblast-like differentiation in a hybrid hydrogel platform, Adv. Healthc. Mater. 4 (2015) 121–130. https://doi.org/10.1002/adhm.201400029.

7. T. Nguyen, K.E. Watkins, V. Kishore, Photochemically crosslinked cell-laden methacrylated collagen hydrogels with high cell viability and functionality, J. Biomed. Mater. Res. A 107 (2019) 1541–1550. https://doi.org/10.1002/jbm.a.36668.

8. K.E. Drzewiecki, J.N. Malavade, I. Ahmed, C.J. Lowe, D.I. Shreiber, A thermoreversible, photocrosslinkable collagen bio-ink for free-form fabrication of scaffolds for regenerative medicine, Technology 05 (2017) 185–195. https://doi.org/10.1142/s2339547817500091.

9. W.T. Brinkman, K. Nagapudi, B.S. Thomas, E.L. Chaikof, Photocrosslinking of type I collagen gels in the presence of smooth muscle cells: Mechanical properties, cell viability, and function, Biomacromolecules 4 (2003) 890–895. https://doi.org/10.1021/bm0257412.

10. E. Nicol, Photopolymerized porous hydrogels, Biomacromolecules 22 (2021) 1325–1345. https://doi.org/10.1021/acs.biomac.0c01671.

11. C. Yu, J. Schimelman, P. Wang, K.L. Miller, X. Ma, S. You, J. Guan, B. Sun, W. Zhu, S. Chen, Photopolymerizable biomaterials and light-based 3D printing strategies for biomedical applications, Chem. Rev. 120 (2020) 10695–10743. https://doi.org/10.1021/acs.chemrev.9b00810.

12. K. Yue, G.T. Santiago, M.M. Alvarez, A. Tamayol, N. Annabi, A. Khademhosseini, Synthesis, properties, and biomedical applications of gelatin methacryloyl (GelMA) hydrogels, Biomaterials 73 (2015) 254–271. https://doi.org/10.1016/j.biomaterials.2015.08.045.

13. A.A. Appel, M.A. Anastasio, J.C. Larson, E.M. Brey, Imaging challenges in biomaterials and tissue engineering, Biomaterials 34 (2013) 6615–6630. https://doi.org/10.1016/j.biomaterials.2013.05.033.

14. S.Y. Nam, L.M. Ricles, L.J. Suggs, S.Y. Emelianov, Imaging strategies for tissue engineering applications, Tissue Eng. B 21 (2015) 88–102. https://doi.org/10.1089/ten.teb.2014.0180.

15. M.A. Rice, K.R. Waters, K.S. Anseth, Ultrasound monitoring of cartilaginous matrix evolution in degradable PEG hydrogels, Acta Biomater. 5 (2009) 152–161. https://doi.org/10.1016/j.actbio.2008.07.036.

16. M. Gudur, R.R. Rao, Y.-S. Hsiao, A.W. Peterson, C.X. Deng, J.P. Stegemann, Noninvasive, quantitative, spatiotemporal characterization of mineralization in three-dimensional collagen hydrogels using high-resolution spectral ultrasound imaging, Tissue Eng. Part C Methods 18 (2012) 935–946. https://doi.org/10.1089/ten.tec.2012.0180.

17. B. Shrestha, K. Stojkova, R. Yi, M.A. Anastasio, J.Y. Ye, E.M. Brey, Gold nanorods enable noninvasive longitudinal monitoring of hydrogels *in vivo* with photoacoustic tomography, Acta Biomater. 117 (2020) 374–383. https://doi.org/10.1016/j.actbio.2020.09.048.

18. M. Zhang, Z. Wang, P. Huang, G. Jiang, C. Xu, W. Zhang, R. Guo, W. Li, X. Zhang, Real-time and noninvasive tracking of injectable hydrogel degradation using functionalized AIE nanoparticles, Nanophotonics 9 (2020) 2063–2075. https://doi.org/10.1515/nanoph-2020-0087.

19. N. Artzi, N. Oliva, C. Puron, S. Shitreet, S. Artzi, A. bon Ramos, A. Groothuis, G. Sahagian, E.R. Edelman, In vivo and in vitro tracking of erosion in biodegradable materials using noninvasive fluorescence imaging, Nat. Mater. 10 (2011) 890–890. https://doi.org/10.1038/nmat3095.

20. W. Wang, J. Liu, C. Li, J. Zhang, J. Liu, A. Dong, D. Kong, Real-time and non-invasive fluorescence tracking of in vivo degradation of the thermosensitive PEGlyated polyester hydrogel, J. Mater. Chem. B. 2 (2014) 4185–4192. https://doi.org/10.1039/c4tb00275j.

21. Z. Cheng, R. Chai, P. Ma, Y. Dai, X. Kang, H. Lian, Z. Hou, C. Li, J. Lin, Multiwalled carbon nanotubes and NaYF4:Yb^3+^/Er^3+^ nanoparticle-doped bilayer hydrogel for concurrent NIR-triggered drug release and up-conversion luminescence tagging, Langmuir 29 (2013) 9573–9580. https://doi.org/10.1021/la402036p.

22. G. Jalani, R. Naccache, D.H. Rosenzweig, S. Lerouge, L. Haglund, F. Vetrone, M. Cerruti, Real-time, non-invasive monitoring of hydrogel degradation using LiYF4 :Yb^3+^/Tm^3+^ NIR-to-NIR upconverting nanoparticles, Nanoscale 7 (2015) 11255–11262. https://doi.org/10.1039/c5nr02482j.

23. Y. Dong, G. Jin, C. Ji, R. He, M. Lin, X. Zhao, A. Li, T.J. Lu, F. Xu, Non-invasive tracking of hydrogel degradation using upconversion nanoparticles, Acta Biomater. 55 (2017) 410–419. https://doi.org/10.1016/j.actbio.2017.04.016.

24. W. Zhu, C. Chu, S. Kuddannaya, Y. Yuan, P. Walczak, A. Singh, X. Song, J.W.M. Bulte, *In vivo* imaging of composite hydrogel scaffold degradation using CEST MRI and two-color NIR imaging, Adv. Funct. Mater. 29 (2019) 1903753. https://doi.org/10.1002/adfm.201903753.

25. M.E. Mertens, A. Hermann, A. Bühren, L. Olde-Damink, D. Möckel, F. Gremse, J. Ehling, F. Kiessling, T. Lammers, Iron oxide-labeled collagen scaffolds for non-invasive MR imaging in tissue engineering, Adv. Funct. Mater. 24 (2013) 754–762. https://doi.org/10.1016/0142-9612(96)81413-x.

26. Z. Chen, C. Yan, S. Yan, Q. Liu, M. Hou, Y. Xu, R. Guo, Non-invasive monitoring of *in vivo* hydrogel degradation and cartilage regeneration by multiparametric MR imaging, Theranostics 8 (2018) 1146–1158. https://doi.org/10.7150/thno.22514.

27. S. Hu, Y. Zhou, Y. Zhao, Y. Xu, F. Zhang, N. Gu, J. Ma, M.A. Reynolds, Y. Xia, H.H.K. Xu, Enhanced bone regeneration and visual monitoring via superparamagnetic iron oxide nanoparticle scaffold in rats, J. Tissue Eng. Regen. Med. 12 (2018) e2085–e2098. https://doi.org/10.1002/term.2641.

28. S.M. Forton, M.T. Latourette, M. Parys, M. Kiupel, D. Shahriari, J.S. Sakamoto, E.M. Shapiro, *In vivo* microcomputed tomography of nanocrystal-doped tissue engineered scaffolds, ACS Biomater. Sci. Eng. 2 (2016) 508–516. https://doi.org/10.1021/acsbiomaterials.5b00476.

29. T.A. Finamore, T.E. Curtis, J.V. Tedesco, K. Grandfield, R.K. Roeder, Nondestructive, longitudinal measurement of collagen scaffold degradation using computed tomography and gold nanoparticles, Nanoscale 11 (2019) 4345–4354. https://doi.org/10.1039/c9nr00313d.

30. S. Uman, L.L. Wang, S.L. Thorn, Z. Liu, J.S. Duncan, A.J. Sinusas, J.A. Burdick, Imaging of Imaging of injectable hydrogels delivered into myocardium with SPECT/CT, Adv. Healthc. Mater. 9 (2020) 2000294. https://doi.org/10.1002/adhm.202000294.

31. L.E. Cole, R.D. Ross, J.M. Tilley, T. Vargo-Gogola, R.K. Roeder, Gold nanoparticles as contrast agents in x-ray imaging and computed tomography, Nanomedicine 10 (2015) 321–341. https://doi.org/10.2217/nnm.14.171.

32. J.C. Hsu, L.M. Nieves, O. Betzer, T. Sadan, P.B. Noel, R. Popovtzer, D.P. Cormode, Nanoparticle contrast agents for X-ray imaging applications, WIREs Nanomed. Nanobiotechnol. (2020) e1642. https://doi.org/10.1002/wnan.1642.

33. P. Thoniyot, M.J. Tan, A.A. Karim, D.J. Young, X.J. Loh, Nanoparticle–hydrogel composites: Concept, design, and applications of these promising, multi-functional materials, Adv. Sci. 2 (2015) 1400010. https://doi.org/10.1002/advs.201400010.

34. M. Yadid, R. Feiner, T. Dvir, Gold nanoparticle-integrated scaffolds for tissue engineering and regenerative medicine, Nano Lett. 19 (2019) 2198–2206. https://doi.org/10.1021/acs.nanolett.9b00472.

35. A.J. Clasky, J.D. Watchorn, P.Z. Chen, F.X. Gu, From prevention to diagnosis and treatment: biomedical applications of metal nanoparticle-hydrogel composites, Acta Biomater. 122 (2021) 1–25. https://doi.org/10.1016/j.actbio.2020.12.030.

36. D.N. Heo, W.-K. Ko, M.S. Bae, J.B. Lee, D.-W. Lee, W. Byun, C.H. Lee, E.-C. Kim, B.-Y. Jung, I.K. Kwon, Enhanced bone regeneration with a gold nanoparticle–hydrogel complex, J. Mater. Chem. B 2 (2014) 1584–1593. https://doi.org/10.1039/c3tb21246g.

37. A. Navaei, H. Saini, W. Christenson, R.T. Sullivan, R. Ros, M. Nikkhah, Gold nanorod-incorporated gelatin-based conductive hydrogels for engineering cardiac tissue constructs, Acta Biomater. 41 (2016) 133–146. https://doi.org/10.1016/j.actbio.2016.05.027.

38. N. Celikkin, S. Mastrogiacomo, X.F. Walboomers, W. Swieszkowski, Enhancing X-ray attenuation of 3D printed gelatin methacrylate (gelMA) hydrogels utilizing gold nanoparticles for bone tissue engineering applications, Polymers 11 (2019) 367. https://doi.org/10.3390/polym11020367.

39. J. Li, D.J. Mooney, Designing hydrogels for controlled drug delivery, Nat. Rev. Mater. 1 (2016) 16071. https://doi.org/10.1038/natrevmats.2016.71.

40. L. Castaneda, J. Valle, N. Yang, S. Pluskat, K. Slowinska, Collagen cross-linking with Au nanoparticles, Biomacromolecules 9 (2008) 3383–3388. https://doi.org/10.1021/bm800793z.

41. T. Schuetz, N. Richmond, M.E. Harmon, J. Schuetz, L. Castaneda, K. Slowinska, The microstructure of collagen type I gel cross-linked with gold nanoparticles, Colloids Surf. B Biointerfaces 101 (2013) 118–125. https://doi.org/10.1016/j.colsurfb.2012.06.006.

42. G. Marcelo, M. López-González, F. Mendicuti, M.P. Tarazona, M. Valiente, Poly(*N*-isopropylacrylamide)/gold hybrid hydrogels prepared by catechol redox chemistry. Characterization and smart tunable catalytic activity, Macromolecules 47 (2014) 6028–6036. https://doi.org/10.1021/ma501214k.

43. D. Lee, D.N. Heo, H.R. Nah, S.J. Lee, W.-K. Ko, J.S. Lee, H.-J. Moon, J.B. Bang, Y.-S. Hwang, R.L. Reis, I.K. Kwon, Injectable hydrogel composite containing modified gold nanoparticles: implication in bone tissue regeneration, Int. J. Nanomed. 13 (2018) 7019–7031. https://doi.org/10.2147/ijn.s185715.

44. R.D. Ross, L.E. Cole, J.M.R. Tilley, R.K. Roeder, Effect of functionalized gold nanoparticle size on X-ray attenuation and substrate binding affinity, Chem. Mater. 26 (2014) 1187–1194. https://doi.org/10.1021/cm4035616.

45. H. Shirahama, B.H. Lee, L.P. Tan, N.-J. Cho, Precise tuning of facile one-pot gelatin methacryloyl (gelMA) synthesis, Sci. Rep. 6 (2016) 31036. https://doi.org/10.1038/srep31036.

46. H. Cui, S. Miao, T. Esworthy, X. Zhou, S. Lee, C. Liu, Z. Yu, J.P. Fisher, M. Mohiuddin, L.G. Zhang, 3D bioprinting for cardiovascular regeneration and pharmacology, Adv. Drug Delivery Rev. 132 (2018) 252–269. https://doi.org/10.1016/j.addr.2018.07.014.

47. D. Loessner, C. Meinert, E. Kaemmerer, L.C. Martine, K. Yue, P.A. Levett, T.J. Klein, F.P.W. Melchels, A. Khademhosseini, D.W. Hutmacher, Functionalization, preparation and use of cell-laden gelatin methacryloyl–based hydrogels as modular tissue culture platforms, Nat. Protoc. 11 (2016) 727–746. https://doi.org/10.1038/nprot.2016.037.

48. K. Rahali, G.B. Messaoud, C.J.F. Kahn, L. Sanchez-Gonzalez, M. Kaci, F. Cleymand, S. Fleutot, M. Linder, S. Desobry, E. Arab-Tehrany, Synthesis and characterization of nanofunctionalized gelatin methacrylate hydrogels, Int. J. Mol. Sci. 18 (2017) 2675. https://doi.org/10.3390/ijms18122675.

49. W.C. Oliver, G.M. Pharr, Measurement of hardness and elastic modulus by instrumented indentation: Advances in understanding and refinements to methodology, J. Mater. Res. 19 (2004) 3–20. https://doi.org/10.1557/jmr.2004.19.1.3.

50. Y.C. Dong, M. Hajfathalian, P.S.N. Maidment, J.C. Hsu, P.C. Naha, S. Si-Mohamed, M. Breuilly, J. Kim, P. Chhour, P. Douek, H.I. Litt, D.P. Cormode, Effect of gold nanoparticle size on their properties as contrast agents for computed tomography, Sci. Rep. 9 (2019) 14912. https://doi.org/10.1038/s41598-019-50332-8.

51. L.E. Marbella, J.E. Millstone, NMR techniques for noble metal nanoparticles, Chem. Mater. 27 (2015) 2721–2739. https://doi.org/10.1021/cm504809c.

52. P.K. Jain, K.S. Lee, I.H. El-Sayed, M.A. El-Sayed, Calculated absorption and scattering properties of gold nanoparticles of different size, shape, and composition: Applications in biological imaging and biomedicine, J. Phys. Chem. B 110 (2006) 7238–7248. https://doi.org/10.1021/jp057170o.

53. B.D. Fairbanks, M.P. Schwartz, C.N. Bowman, K.S. Anseth, Photoinitiated polymerization of PEG-diacrylate with lithium phenyl-2,4,6-trimethylbenzoylphosphinate: polymerization rate and cytocompatibility, Biomaterials 30 (2009) 6702–6707. https://doi.org/10.1016/j.biomaterials.2009.08.055.

54. W. Krause, Delivery of diagnostic agents in computed tomography, Adv. Drug Delivery Rev. 37 (1999) 159–173. https://doi.org/10.1016/s0169-409x(98)00105-7.

55. L. Ouyang, J.P. Armstrong, Y. Lin, J.P. Wojciechowski, C. Lee-Reeves, D. Hachim, K. Zhou, J.A. Burdick, M.M. Stevens, Expanding and optimizing 3D bioprinting capabilities using complementary network bioinks, Sci. Adv. 6 (2020) eabc5529. https://doi.org/10.1126/sciadv.abc5529

56. X. Bai, M. Gao, S. Syed, J. Zhuang, X. Xu, X.Q. Zhang, Bioactive hydrogels for bone regeneration, Bioactive Mater. 3 (2018) 401–417. https://doi.org/10.1016/j.bioactmat.2018.05.006

57. Y. Piao, H. You, T. Xu, H.P. Bei, I.Z. Piwko, Y.Y. Kwan, X. Zhao, Biomedical applications of gelatin methacryloyl hydrogels. Eng. Regen. 2 (2021) 47–56. https://doi.org/10.1016/j.engreg.2021.03.002

58. K. Zhu, S.R. Shin, T. van Kempen, Y. Li, V. Ponraj, A. Nasajpour, S. Mandla, N. Hu, X. Liu, J. Leijten, Y. Lin, M.A. Hussain, Y.S. Zhang, A. Tamayol, A. Khademhosseini, Gold nanocomposite bioink for printing 3D cardiac constructs, Adv. Funct. Mater. 27 (2017) 1605352. https://doi.org/10.1002/adfm.201605352

59. Q. Li, B. Zhang, Q. Xue, C. Zhao, Y. Luo, H. Zhou, L. Ma, H. Yang, D. Bai, A systematic thermal analysis for accurately predicting the extrusion printability of alginate-gelatin-based hydrogel bioinks, Int. J. Bioprinting 7 (2021) 394. https://doi.org/10.18063/ijb.v7i3.394

60. A.D. Cetnar, M.L. Tomov, L. Ning, B. Jing, A.S. Theus, A. Kumar, A.N. Wijntjes, S.R. Bhamidipati, K.P. Do, A. Mantalaris, J.N. Oshinski, R. Avazmohammadi, B.D. Lindsey, H.D. Bauser-Heaton, V. Serpooshan, Patient-specific 3D bioprinted models of developing human heart, Adv. Healthc. Mater. 10 (2021) e2001169. https://doi.org/10.1002/adhm.202001169

61. L. Ning, N. Betancourt, D.J. Schreyer, X. Chen, Characterization of cell damage and proliferative ability during and after bioprinting, ACS Biomater Sci. Eng. 12 (2018) 3906–3918. https://doi.org/10.1021/acsbiomaterials.8b00714

62. L. Ning, B. Yang, F. Mohabatpour, N. Betancourt, M.D. Sarker, P. Papagerakis, X. Chen, Process-induced cell damage: pneumatic versus screw-driven bioprinting, Biofabrication 12 (2020) 025011. https://doi.org/10.1088/1758-5090/ab5f53

63. J. Yin, M. Yan, Y. Wang, J. Fu, H. Suo, 3D bioprinting of low-concentration cell-laden gelatin methacrylate (GelMA) bioinks with a two-step cross-linking strategy, ACS Appl. Mater. Interfaces 10 (2018) 6849–6857. https://doi.org/10.1021/acsami.7b16059

