## Supplementary Data for "Methacrylate-Modified Gold Nanoparticles Enable Non-Invasive Monitoring of Photocrosslinked Hydrogel Scaffolds"

**Table S1.** The effect of Au NP concentration and UV light intensity on photocrosslinking gelMA+Au NP hydrogels. Prepolymer solutions were incubated with photoinitiator for 1 h prior to photocrosslinking.

| [GelMA] (w/v) | [Irgacure 2959] (w/v) | [Au] (mM) | UV intensity (mW/cm$^2$) | Irradiation Time (min) | Photocrosslinking |
|---|---|---|---|---|---|
| 10% | 0.5% | 0 | 7 | 6 | ++ |
| | | 3.0 | 7 | 6 | ++ |
| | | 7.4 | 7 | 6 | ++ |
| | | 7.4 | 15 | 6 | ++ |
| | | 14.9 | 15 | 6 | - |
| | | 22.3 | 15 | 6 | - |

++ fully crosslinked, + partially crosslinked, - little or no crosslinking observed.

**Table S2.** The effect of the Au NP concentration, photoinitiator (Irgacure 2959) concentration, UV light intensity, and irradiation time on photocrosslinking gelMA+Au NP hydrogels comprising 10% w/v gelMA. Prepolymer solutions were incubated with photoinitiator for 1 h prior to photocrosslinking.

| [Irgacure] (w/v) | [Au] (mM) | UV intensity, irradiation time | | |
|---|---|---|---|---|
| | | 7 mW/cm$^2$, 6 min | 15 mW/cm$^2$, 6 min | 30 mW/cm$^2$, 4 min |
| 0.5% | 7 | ++ | ++ | ++ |
| | 10 | + | + | + |
| | 15 | - | - | - |
| 1.0% | 7 | ++ | ++ | ++ |
| | 10 | + | + | + |
| | 15 | - | - | - |

++ fully crosslinked, + partially crosslinked, - little or no crosslinking observed.

**Table S3.** The effect of the Au NP and gelMA concentration on photocrosslinking 1-step gelMA-Au NP hydrogels with 0.5% w/v Irgacure photoinitiator under UV irradiation at 30 mW/cm$^2$ for 6 min. Prepolymer solutions were incubated with photoinitiator for 1 h prior to photocrosslinking.

| [Au] (mM) | [GelMA] | |
|---|---|---|
| | 10% | 20% |
| 15 | - | + |
| 37 | - | + |

++ fully crosslinked, + partially crosslinked, - little or no crosslinking observed.

**Table S4.** The effect of the Au NP concentration, Irgacure or LAP photoinitiator, and incubation time on photocrosslinking 1-step gelMA-Au NP hydrogels with 20% w/v gelMA and 0.5% w/v photoinitiator under UV irradiation at 30 mW/cm$^2$ for 4 min respectively.

| [Au] mM | Irgacure 2959 | | LAP | | |
|---|---|---|---|---|---|
| | 1 h | 24 h | 1 h | 24 h | 7 d |
| 0 | ++ | ++ | ++ | ++ | ++ |
| 4.4 | ++ | ++ | ++ | ++ | ++ |
| 10 | + | + | + | ++ | ++ |
| 15 | - | - | - | + | ++ |
| 22 | - | - | - | + | ++ |
| 37 | - | - | - | + | ++ |

++ fully crosslinked, + partially crosslinked, - little or no crosslinking observed.

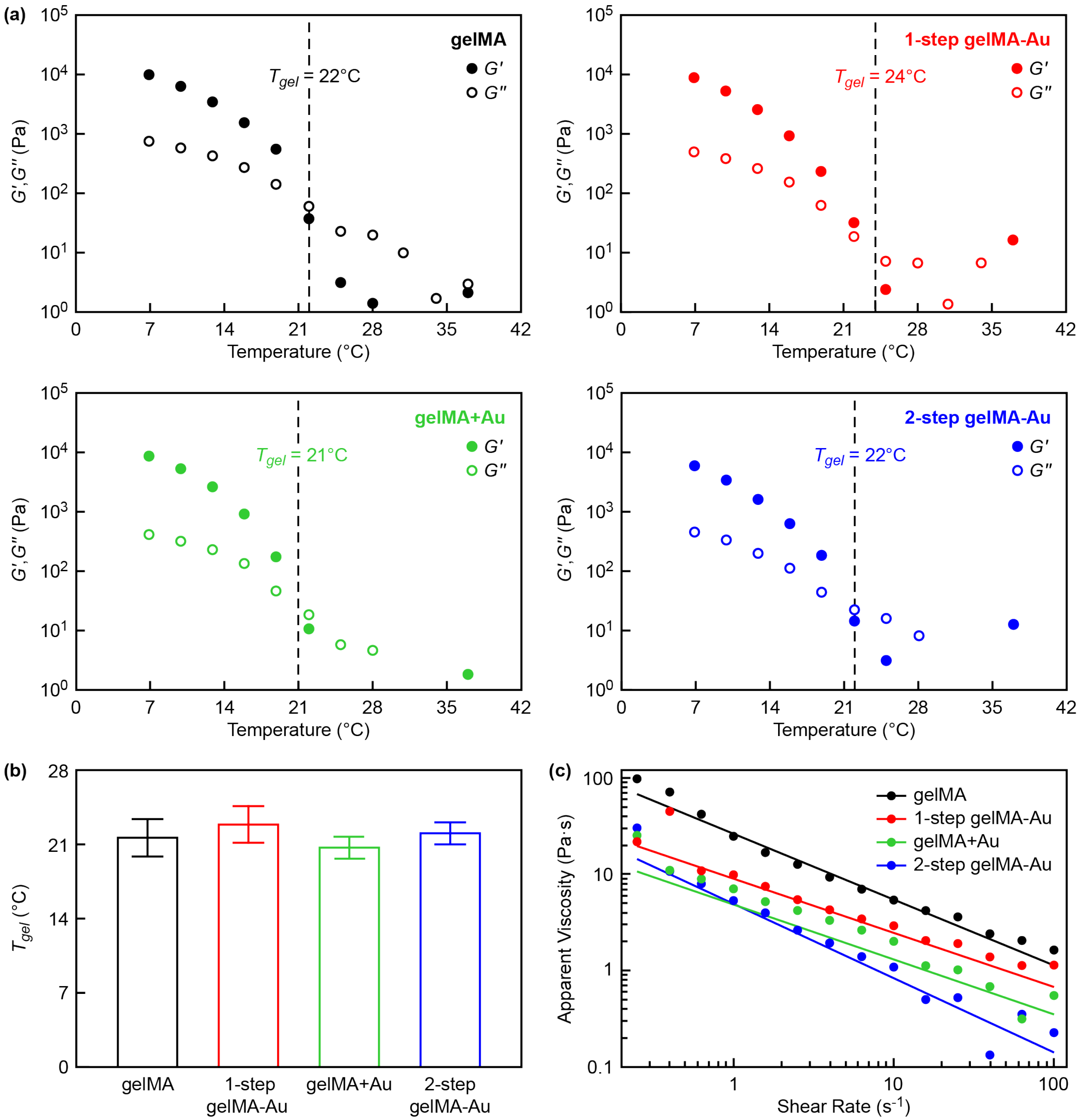


**Fig. S1. Rheology of gelMA and gelMA-Au NP prepolymer solutions.** (a) The sol-gel transition temperature ($T_{gel}$) was measured from the change in storage moduli ($G'$) and loss moduli ($G''$) as a function of temperature during cooling. Data is shown for samples exhibiting the median $T_{gel}$ for each group. (b) Differences in the measured $T_{gel}$ between prepolymer solutions were not statistically significant ($p > 0.23$, ANOVA). Error bars show one standard deviation of the mean ($n = 3$/group). (c) Flow behavior measured by a steady state strain rate sweep indicated that the apparent viscosity of gelMA was greater than 1-step gelMA-Au, and 1-step gelMA-Au was greater than both gelMA+Au and 2-step gelMA-Au ($p < 0.05$, Tukey). Differences between gelMA+Au and 2-step gelMA-Au were not statistically significant. Data points show mean values ($n = 3$/group) and error bars are not shown for clarity.

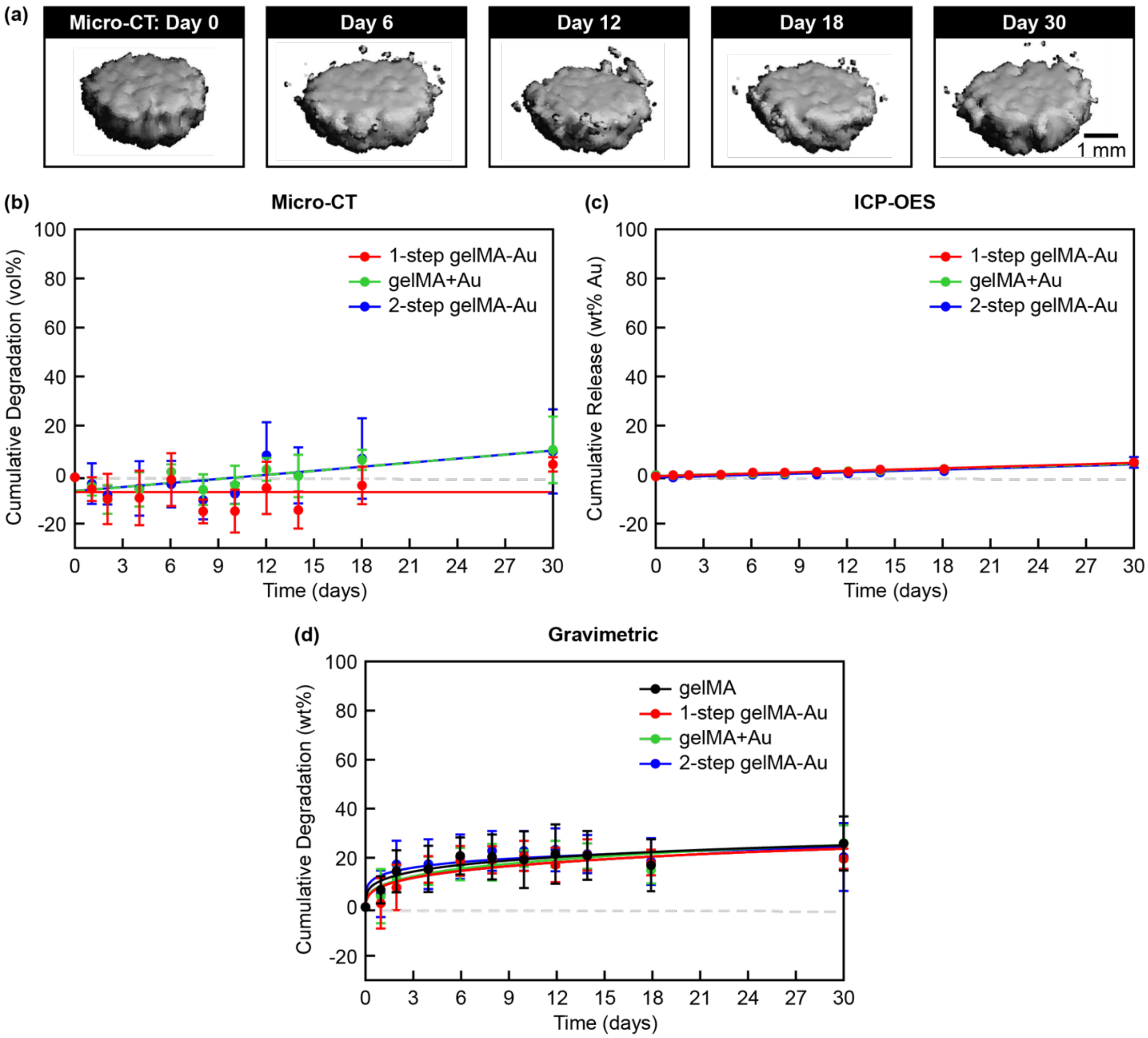


**Fig. S2. Stability of gelMA and gelMA-Au NP hydrogels against *in vitro* hydrolytic degradation.** (a) Representative segmented micro-CT image reconstructions for selected time points showing no apparent change in the volume of 1-step gelMA-Au hydrogels. Degradation kinetics were measured longitudinally by (b) the cumulative change in segmented hydrogel volume using contrast-enhanced micro-CT, (c) the cumulative release of Au NPs into the media using ICP-OES, and (d) the cumulative hydrogel mass loss using gravimetric analysis. All gelMA and gelMA-Au NP hydrogels were stable against hydrolysis for at least one month. Differences between gelMA and gelMA-Au NP hydrogels were not statistically significant. Degradation kinetics measured by micro-CT and ICP-OES were modeled by linear least squares regression. Degradation kinetics measured by gravimetrical analysis were modeled by non-linear least squares regression using a four-parameter logistic model (Eq. 1) with a log-transform of time. Error bars show one standard deviation of the mean ($n$ = 5/group). Error bars not shown lie within the data point.

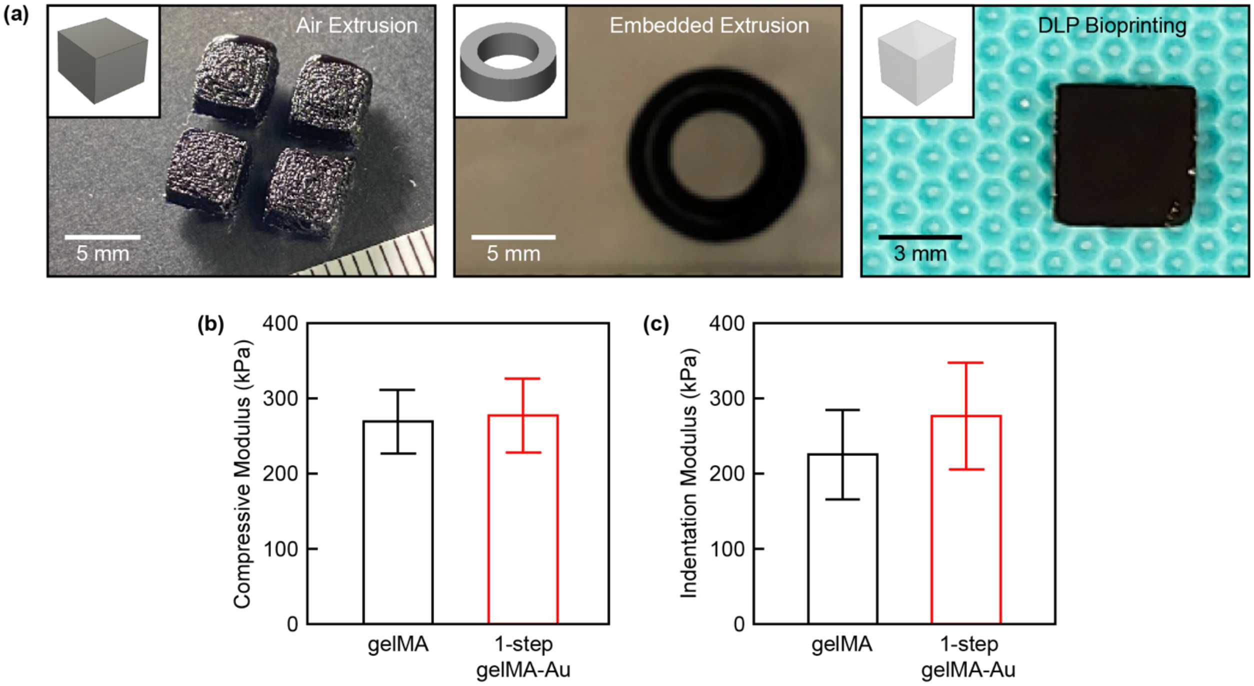


**Fig. S3. Mechanical properties of 3D bioprinted 1-step gelMA-Au hydrogel constructs.** (a) Cubic scaffolds printed by air extrusion, a cylindrical tube mimicking a blood vessel printed by embedded extrusion, and a cubic scaffold printed by DLP bioprinting. Insets show corresponding CAD models. The (b) compressive modulus and (c) microindentation modulus of cubic scaffolds printed by air extrusion and measured after reaching equilibrium swelling. Differences between gelMA and 1-step gelMA-Au NP hydrogels were not statistically significant. Error bars show one standard deviation of the mean ($n$ = 3-4/group).

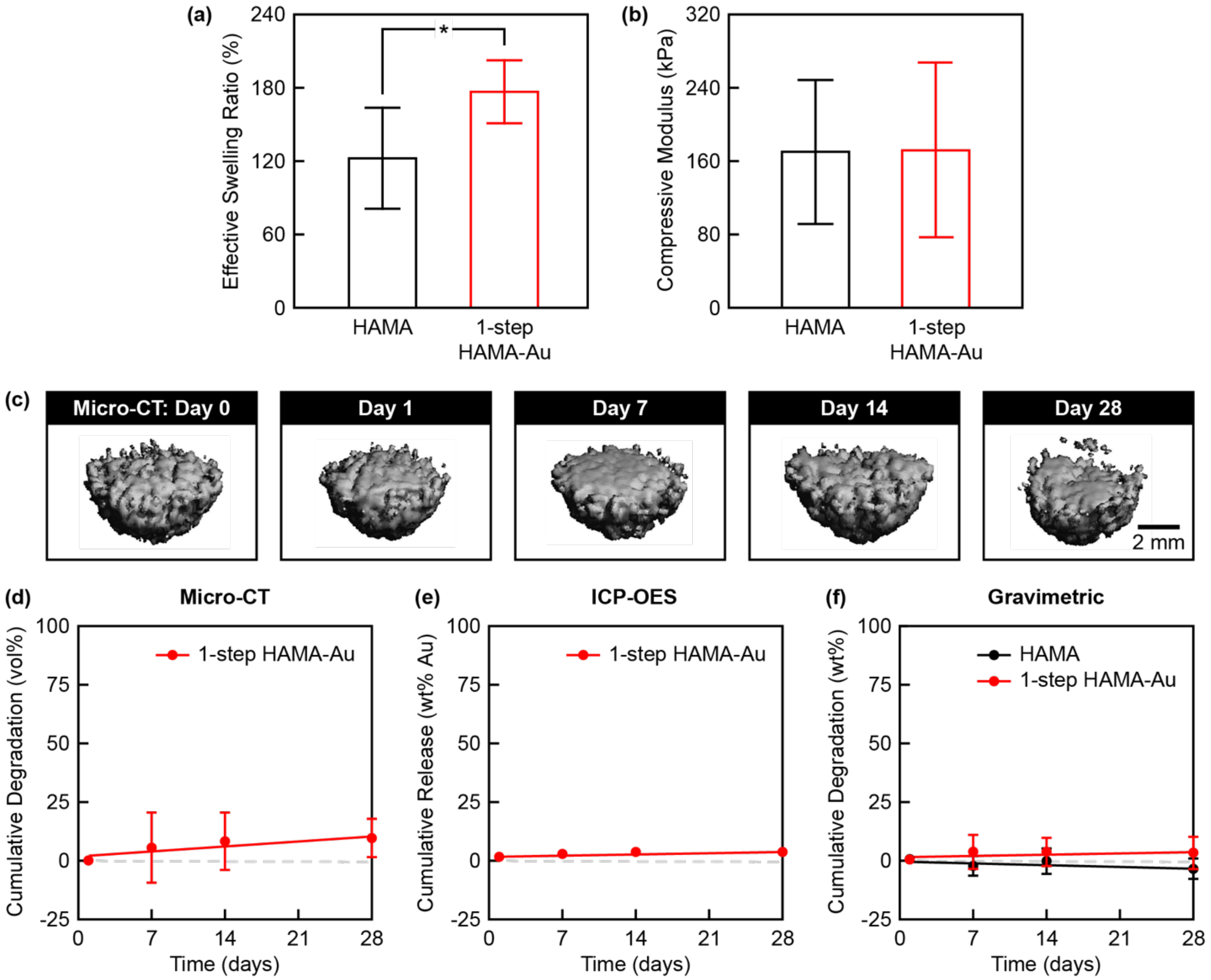


**Fig. S4. Characterization of HAMA and 1-step HAMA-Au NP hydrogels.** The (a) effective swelling ratio ($n$ = 7/group) and (b) compressive modulus ($n$ = 4-6/group) of HAMA and 1-step HAMA-Au NP hydrogels after reaching equilibrium swelling. The effective swelling ratio of 1-step HAMA-Au hydrogels was greater than HAMA alone, but the compressive modulus was not statistically different. Error bars show one standard deviation of the mean. *$p < 0.05$, Tukey. (c) (c) Representative segmented micro-CT image reconstructions for selected time points showing no apparent volume loss of 1-step HAMA-Au hydrogels during *in vitro* hydrolysis for up to four weeks. Degradation kinetics were measured longitudinally by (d) the cumulative change in segmented hydrogel volume using contrast-enhanced micro-CT, (e) the cumulative release of Au NPs into the media using ICP-OES, and (f) the cumulative hydrogel mass loss using gravimetric analysis. Both HAMA and 1-step HAMA-Au NP hydrogels were stable against hydrolysis for at least four weeks. Differences between HAMA and 1-step HAMA-Au NP hydrogels were not statistically significant. Degradation kinetics measured by micro-CT, ICP-OES and gravimetrical analysis were modeled by linear least squares regression. Error bars show one standard deviation of the mean ($n$ = 5/group for micro-CT, $n$ = 7/group for ICP-OES and gravimetric analysis). Error bars not shown lie within the data point.
